# Genetic population structure of Japanese freshwater crab, *Geothelphusa dehaani* species complex (Decapoda: Potamidae) using genome-wide SNP

**DOI:** 10.1101/2025.03.05.641571

**Authors:** Taiga Kunishima, Kento Takata, Kanto Nishikawa, Kodai Gibu, Miyuki Nishijima, Akira Iguchi

## Abstract

The Japanese freshwater crab *Geothelphusa dehaani* species complex is distributed widely across the Japanese Archipelago. Despite its suggested high genetic and morphological diversity, key aspects such as nuclear DNA (nuDNA) population structure and relationship between body color patterns and genetic populations remain unclear. To address these gaps, this study analyzed genome-wide single nucleotide polymorphisms (SNPs) in nuDNA and mitochondrial DNA (mtDNA) cytochrome oxidase subunit 1 (COI) markers in samples from Hokkaido to the Tokara Islands, Japan. Admixture analysis identified five distinct populations with significant geographic boundaries. These populations exhibited unique geographical patterns, spanning across islands and enclave distribution, indicating that *G*. *dehaani* populations have been shaped by complex factors, including sea level changes and volcanic activity. Regional body color variations partially aligned with SNP clades. Further, combining body color with collection locality data could help identify the specimen populations. Contrasting patterns between mtDNA and nuDNA suggest historical gene flow and adaptive introgression, emphasizing the need for caution when interpreting earlier phylogenetic studies based on combined mtDNA and nuDNA sequences. Our findings provide a foundational baseline for future research into the taxonomy, phylogeny, and population dynamics of the *G*. *dehaani* species complex, advancing our understanding of its evolutionary history.

## Introduction

The genetic population structure of a species is shaped by interactions between geological events, climatic changes, and ecological dynamics [1–3]. These processes influence population divergence, gene flow, and adaptation, contributing to current biodiversity. Primary freshwater crabs exhibit a unique evolutionary history, characterized by a land-locked life history and direct development without a planktonic larval stage, and these traits often lead to genetic differentiation and speciation among islands [4–6]. This relationship between life history traits and genetic structure, shaped by the geographical history, makes these crabs a valuable model for understanding genetic divergence mechanisms.

In particular, the genus *Geothelphusa* is the northernmost and diverse freshwater crab genus, spanning from Taiwan to the Japanese Archipelago [6], exhibiting island-specific differentiation in some species [7–9]. *Geothelphusa dehaani* (Japanese name, Sawagani), the most commonly observed freshwater crab in Japan, is usually found in freshwater habitats from coastal areas to high-altitude mountains, from Nakanoshima in the northern Ryukyu Islands (29.9°N) to southern Hokkaido (41.5°N) [9–12]. This wide distribution makes it ideal for investigating how the geological history of the Japanese Archipelago has shaped the genetic structures of these freshwater crab.

Previous genetic studies on *G*. *dehaani* were limited in scope, using samples from restricted areas and examining partial regions of enzyme or mitochondrial DNA sequences [13–16]. Takenaka et al. [17] was the first to comprehensively analyze its phylogeny, revealing 10 populations across the Japanese Islands and identifying island-specific genetic differentiation using a combination of partial mitochondrial DNA (mtDNA) markers, namely cytochrome oxidase subunit 1 (COI) and 16S rRNA and nuclear DNA (nuDNA), namely internal transcribed spacer (ITS) and histone H3. However, the discordance between mtDNA and nuDNA phylogenies, often observed in other taxa, e.g., the geotrupid dung beetle *Phelotrupes auratus* [18] or the Japanese fire-bellied newt *Cynops pyrrhogaster* [19], indicates the need for further analysis. Such discordance can result from differing evolutionary rates or adaptive mtDNA introgression [20]. If this applies to *G*. *dehaani*, it is necessary to examine each genetic structure in detail to understand its phylogeny and evolutionary processes.

Differences in genetic structure often reflect phenotypic traits, such as body coloration. Geographic variation in body color is well-documented in *G*. *dehaani* [15, 21–24], with individuals traditionally classified into three main color types: dark (DA), red (RE), and blue (BL) [21, 22]. These body color types exhibit distinct geographic distributions, with DA being widespread and RE and BL more localized [21–25]. Body color could serve as a population marker if there were a relationship between body color and genetic structure; in this case, *G*. *dehaani* could be used as a model for studying gene flow, selection, and demographic history underlying color variation, as animal coloration is a key adaptive trait with diverse functions [18, 26]. However, previous studies on color– genetic relationships yielded conflicting results, largely due to reliance on partial mtDNA data [14, 15, 17, 24, 27]. As mtDNA data alone offers limited insights into the demographic history and geographic division of color forms, genome-wide sequence data are necessary to elucidate the genetic basis of geographic color variation [18].

Genome-wide approaches using large datasets of single nucleotide polymorphisms (SNPs) in nuDNA help uncovering insights not visible by analyzing a region of mtDNA, nuDNA, or morphological analyses [28]. Further, genome-wide approaches provide a fundamental baseline to tackle taxonomic issues in the *G*. *dehaani* species complex, which may include cryptic species or distinct genetic groups. Molecular phylogenetic studies based on partial mtDNA regions have suggested multiple clades and potential undescribed species within *G*. *dehaani* [14, 15, 17, 24, 27]. Recently, Naruse and Ng [9] described two new species, *G*. *mutsu* and *G*. *amakusa*, from Aomori in Honshu Island and Amakusa Island, respectively, redesignating the lectotype and type locality of *G*. *dehaani*. However, Takenaka et al. [17] found no genetic differences between specimens from Amakusa in the Nagasaki Prefecture and those near the *G. dehaani* type locality (Nagasaki City, Clade 7). Furthermore, no phylogenetic relationship was described for these two species by Naruse and Ng [9], indicating a complex phylogenetic relationship of the *G. dehaani* species complex, including *G*. *amakusa* and *G*. *mutsu*.

Addressing these issues requires extensive sampling across the *G. dehaani* species complex distribution and advanced molecular tools, such as next-generation sequencing [17, 27]. Accordingly, this study aimed to investigate the genetic population structure of the *G. dehaani* species complex using genome-wide SNP data. We comprehensively collected 1,039 specimens from 299 sites across the Japanese Islands to examine the phylogenetic relationships between nuDNA and mtDNA and assess a potential discordance. Moreover, we used approximate Bayesian computational (ABC) analysis to infer its demographic history and analyzed the relationship between body color and genetic structure to evaluate its potential as a population marker. Finally, we have discussed the evolutionary patterns of the *G. dehaani* species complex in the Japanese Islands.

## Results

### Body color distributions

In this study, we identified five body color types: DA, RE, BL, amakusa, and other color (OC). The OC type, which did not align with descriptions by Chokki [22] and Naruse and Ng [9], featured a reddish purple carapace with a milky white wedge pattern in the posterior two-thirds or a dark brown anterior half with a dark orange pattern on the anterior margin and posterior half, all with whitish bases on the thoracic legs (Fig. 1, Table S1). As expected, the distribution patterns in the Japanese Archipelago differed among body color types (Fig. 1): DA was widespread in Hokkaido, Honshu, Shikoku, Kyushu, and adjacent islands (Sadogashima, Okinoshima, and Goto islands); BL was locally concentrated along the Pacific coast, particularly in Honshu from the Boso Peninsula to the Izu Peninsula and the southeastern Kii Peninsula, southern part of Shikoku, southern-west part of Kyushu, and Osumi Islands (Yakushima and Tanegashima islands); RE was primarily found in southern Japan, including the southern Kii Peninsula, southwestern Shikoku, and Kyushu; amakusa was located in the Amakusa Islands; and OC in Nagasaki and Oita in Kyushu.

**Figure 1.**
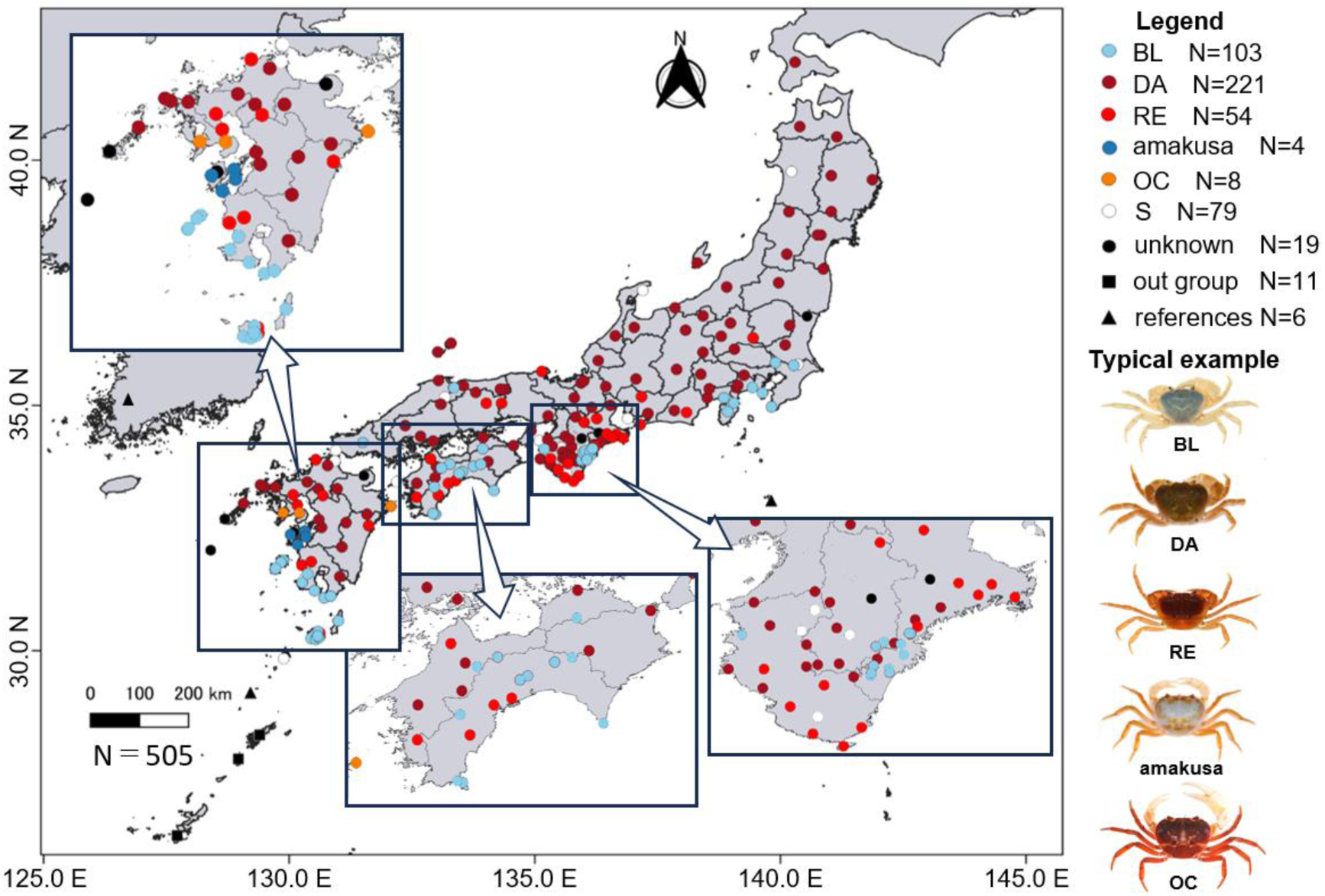
Map showing the geographical distribution of the analyzed samples. Symbol shapes represent species and data sources; circles, the *Geothelphusa dehaani* species complex; squares, outgroups; triangles: reference data of *G*. *dehaani* sensu lato from the literature. Symbol colors represent body coloration types; sky blue, BL type; brown, DA type; red, RE type; blue, amakusa type: orange, OC type; white, unidentified specimens due to small individuals; and black, unknown.

### Genetic population structure based on mtDNA

The amplified mtDNA sequences ranged from 557 to 590 base pairs, yielding 195 haplotypes (Table S2). The *G. dehaani* species complex was divided into three monophyletic populations (clades pop1–3) and 14 subclades in the neighbor network (Fig. 2). Pop1 comprised specimens from Shikoku and the southeastern Kii Peninsula, further divided into three subpopulations: 1a (central and western Shikoku), 1b (southeastern Kinki), and 1c (southeastern Shikoku). Pop2 included specimens from Shikoku and Kyushu, comprising 2a (Kyushu) and 2b (Shikoku and Kyushu). Pop3 was separated into nine subpopulations: 3a (Hokkaido, Honshu from Tohoku to eastern Chugoku, and eastern Shikoku), 3b (western Chugoku and northwestern Shikoku), 3c (western Kyushu, including *G. amakusa*, and South Korea), 3d (northeastern Shikoku and Kyushu), 3e (Yakushima and Nakanoshima islands), 3f (Tanegashima Islands), 3g (Koshikishima Islands), 3h (southern Kanto and Eastern Shizuoka Prefecture), and 3i (Danjo Islands in Oshima Islands).

**Figure 2.**
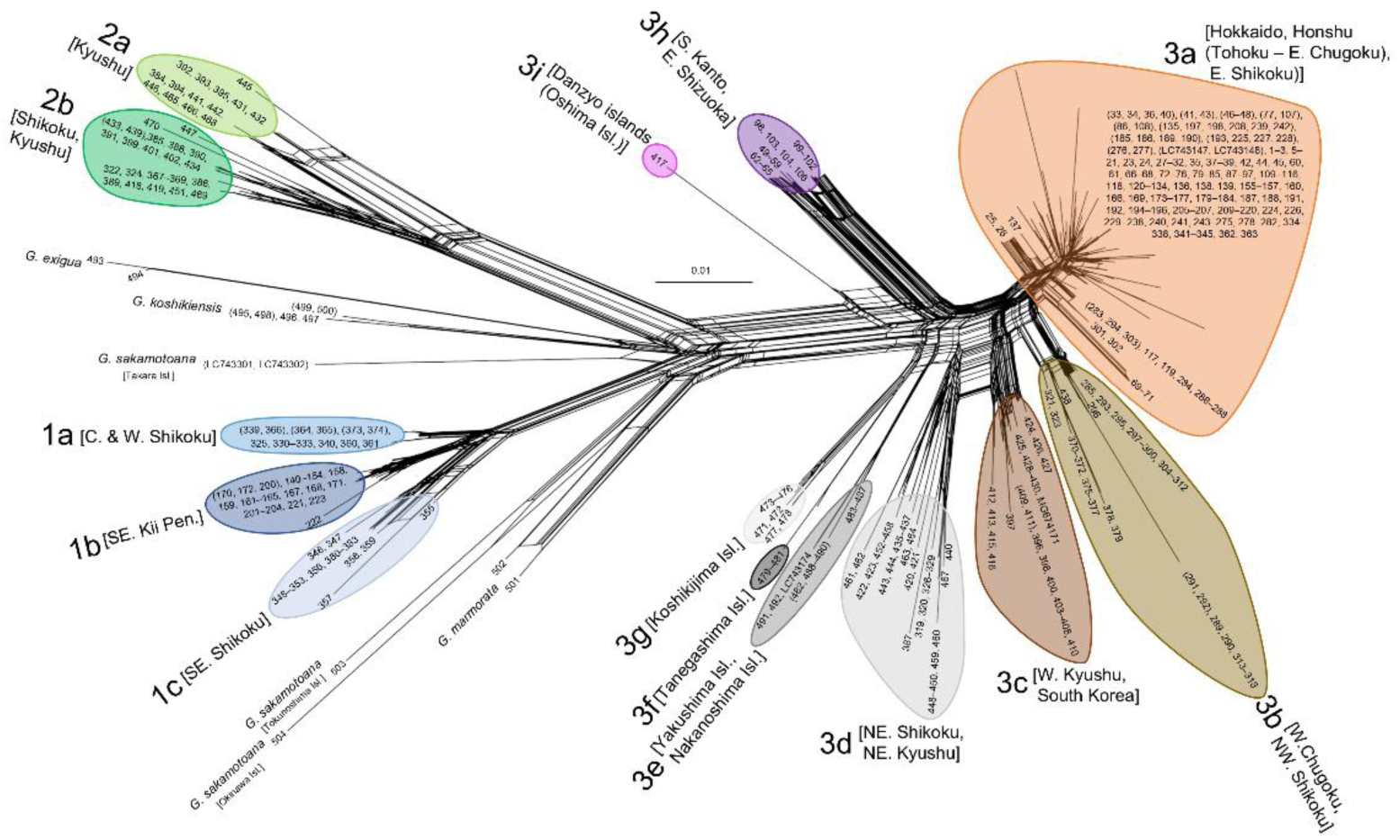
Neighbor-net phylogenetic network based on the 557–590 base pairs of the mtDNA COI region.

The haplotype diversity for pop1, pop2, and pop3 were 0.870, 0.955, and 0.970, respectively, with a nucleotide diversity of 1.66%, 2.73%, and 2.32%, respectively (Table S3). Significant Tajima’s *D* and Fu’s *Fs* values were obtained only for pop3 (*p* < 0.01), with values of −1.68 (*p* = 0.009) and −23.6 (*p* = 0.006), respectively. Pairwise fixation index (*F*_ST_) values were 0.731 between pop1 and pop2, 0.749 between pop1 and pop3, and 0.752 between pop2 and pop3, with *p* = 0 for all sites (Table S4).

### Genetic differences among populations based on MIG-seq

Table S5 showed the number of SNPs and CV error values for datasets 0–6. For set 0, the lowest CV error value was obtained for K = 5, dividing the population into five groups: HO (Hokkaido, Honshu from Tohoku to eastern Chugoku, and Shikoku), SHI (southeast Kinki and Shikoku), nKC (western Chugoku and northern Kyushu), cK (central Kyushu, including *G*. *amakusa*), and sKK (southern Kyushu, southern Kanto, and eastern Shizuoka) (Fig. 3; Fig. S1). These groups partially corresponded to our observed color types, with HO primarily comprising DA and RE; SHI primarily comprising BL; nKC primarily comprising DA and RE; cK primarily comprising DA, RE, and amakusa; and sKK predominantly comprising BL (Figs. 1 and 3). In the SHI population, only four specimens corresponded to DA and RE types (specimen numbers 62, 276, 432, and 527). A few OC specimens were observed in nKC and cK populations. Among the 140 samples in sets 1–6, including duplicates, 133 had a Q value (Q) of >90% (Fig. S2). Seven samples had Q values between 10% and 90%, including set 1 (No. 104), set 2 (No. 166), set 4 (No. 346), and set 6 (Nos. 457, No. 458, No. 465, No. 483) (Fig. S2). Except for No. 483, these samples were collected near geographical population boundaries.

**Figure 3.**
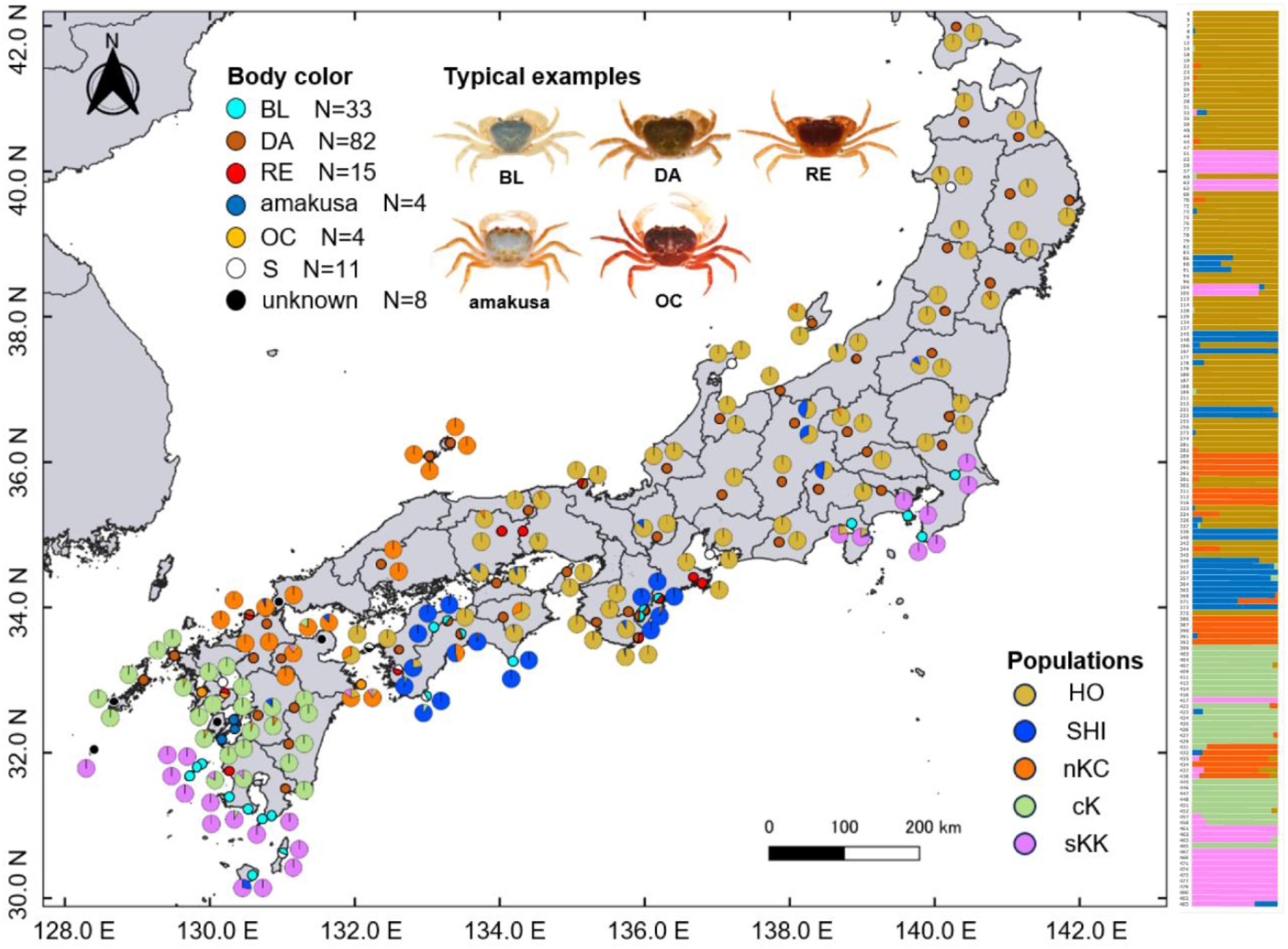
Map showing the results of the admixture analysis for each location (set 0) and the distribution of color types. Large pie chart, genetic elements of each individual; small pie chart, color types of individuals at each location used in this analysis. Color types are indicated as in Figure 1. The colors in the pie charts corresponding to each group and genetic elements are as follows: HO (brown), Hokkaido, Honshu from Tohoku to eastern Chugoku, Shikoku; SHI (blue), southeast Kinki and Shikoku; nKC (orange), western Chugoku and northern Kyushu; cK (green), central Kyushu, including *Geothelphusa amakusa*; and sKK (pink), southern Kyushu, southern Kanto and eastern Shizuoka. The colors of the admixture graph correspond to each genetic element on the map, and the sample numbers refer to Table S1 and Fig. S1.

The results for both variant positions and all positions were consistent in the five populations (Table 1). The highest nucleotide diversity (π) value was observed in sKK, whereas HO and SHI had the lowest. nKC had the highest observed heterozygosity (*Ho*), and sKK the highest expected heterozygosity (*He*). All groups exhibited higher *He* than *Ho*, indicating homozygous excess, with all inbreeding coefficient (*F*_IS_) > 0. Pairwise *F*_ST_ and *F*_ST_ correction value (*F*′_ST_) values indicated a deep genetic divergence for SHI. These values were consistent across populations, being the largest between SHI and cK and smallest between HO and nKC (Table 2).

**Table 1.**
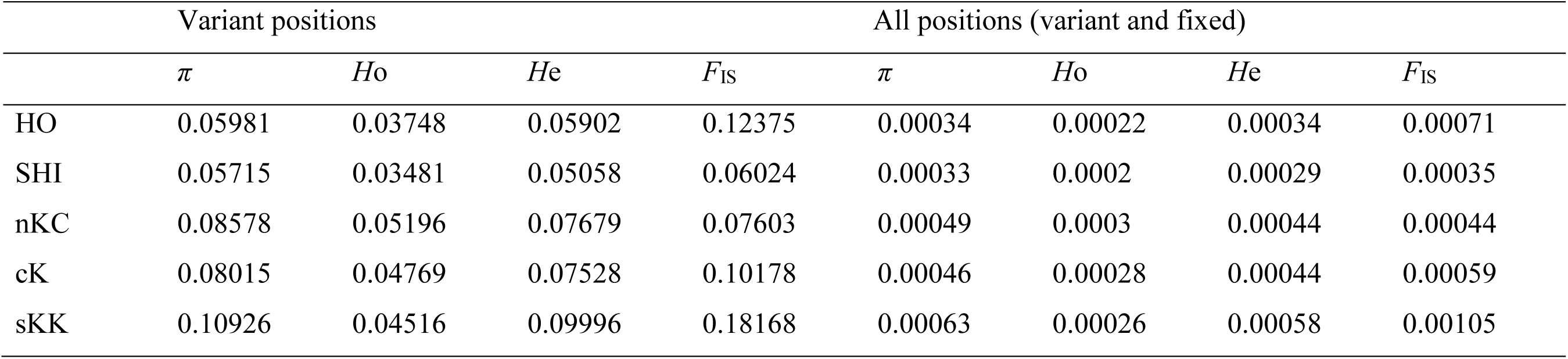
Overall population genomic statistics for the *Geothelphusa dehaani* species complex based on 354 SNP loci (variant positions) and 9058 SNP loci (all positions). Abbreviations: Observed heterozygosity (*Ho*), Expected heterozygosity (*He*), Nucleotide diversity (π), Inbreeding coefficient (*F*_IS_).

|  | Variant positions |  |  |  | All positions (variant and fixed) |  |  |  |
| --- | --- | --- | --- | --- | --- | --- | --- | --- |
| | $\pi$ | *Ho* | *He* | $F_{IS}$ | $\pi$ | *Ho* | *He* | $F_{IS}$ |
| HO | 0.05981 | 0.03748 | 0.05902 | 0.12375 | 0.00034 | 0.00022 | 0.00034 | 0.00071 |
| SHI | 0.05715 | 0.03481 | 0.05058 | 0.06024 | 0.00033 | 0.0002 | 0.00029 | 0.00035 |
| nKC | 0.08578 | 0.05196 | 0.07679 | 0.07603 | 0.00049 | 0.0003 | 0.00044 | 0.00044 |
| cK | 0.08015 | 0.04769 | 0.07528 | 0.10178 | 0.00046 | 0.00028 | 0.00044 | 0.00059 |
| sKK | 0.10926 | 0.04516 | 0.09996 | 0.18168 | 0.00063 | 0.00026 | 0.00058 | 0.00105 |

**Table 2.** *F*_ST_ and *F*′_ST_ values among the five populations of the *Geothelphusa dehaani* species complex based on 354 SNP loci. Bottom left: Pairwise fixation index (*F*_ST_), top right: *F*_ST_ correction value (*F*′_ST_).

| $F_{ST} / F'_{ST}$ | HO | SHI | nKC | cK | sKK |
| --- | --- | --- | --- | --- | --- |
| HO |  | 0.15208 | 0.11779 | 0.1549 | 0.17581 |
| SHI | 0.13479 |  | 0.22696 | 0.21907 | 0.18789 |
| nKC | 0.11826 | 0.19693 |  | 0.16267 | 0.19927 |
| cK | 0.15486 | 0.20279 | 0.14696 |  | 0.15475 |
| sKK | 0.18086 | 0.15804 | 0.18764 | 0.14527 |  |

### Demographic history

To explore demographic history of the five genetic clades, we analyzed representative individuals from a single ancestral population using DIYABC-RF based on 313 SNPs. Principal component analysis (PCA) pre-scenario checks indicated that the observed dataset aligned well with the simulated dataset (Fig. 4**a**), suggesting that the analysis conditions were suitable for random forest analysis.

**Figure 4.**
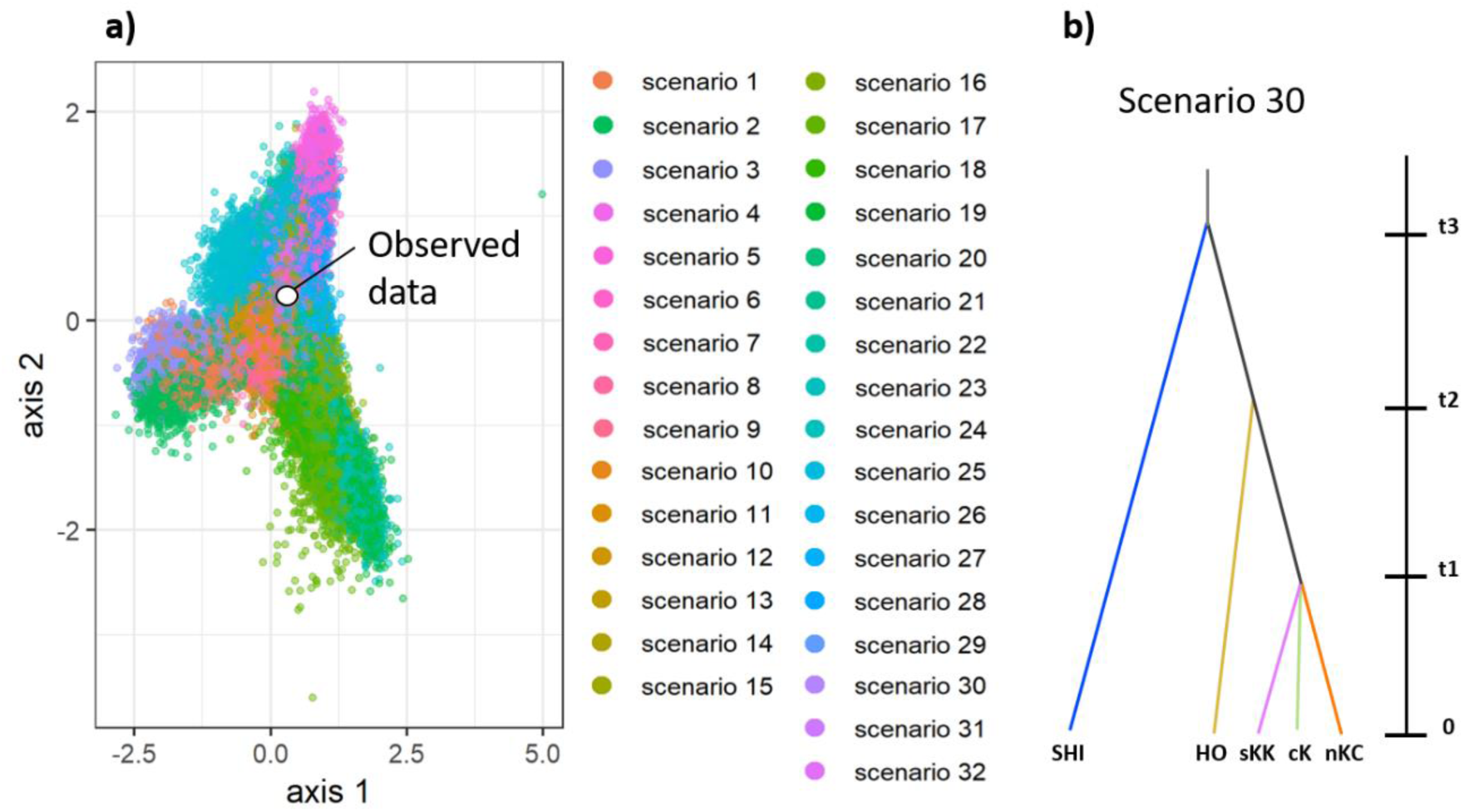
Results of DIYABC-RF for the *Geothelphusa dehaani* species complex based on 313 SNPs. (**a**) Principal component analysis plots evaluating the fit between the observed data and simulated datasets for 32 demographic scenarios of the *G. dehaani* species complex in the Japanese Islands. (**b**) The best voted scenario for the divergence history of *G*. *dehaani* populations (Scenario 30; see Fig. S3).

Among the 32 hypothetical divergence scenarios tested (Fig. S3), Scenario 30 was selected by 52.0% votes (Table S6, 520 out of a total 1000 votes) as the best fit, with a mean posterior probability of 0.831 and prior and posterior error rates (i.e., global and local errors) of 0.133 and 0.169, respectively. Scenario 30 proposed a divergent history where the SHI population diverged first, followed by HO, with nKC, sKK, and cK populations differentiating around the same time (Fig. 4**b**). Scenarios diverging by body color (Scenarios 1–3) received only two votes.

## Discussion

### Genetic population structure and evolutionary history

Our findings indicate that the genetic population structure of the *G. dehaani* species complex was shaped by multiple geological events throughout its complex evolutionary history, marked by repeated population expansions and contractions. Admixture analysis identified five populations across the Japanese Archipelago, delineated by distinct geographical boundaries. Notably, most populations, except the cK population, span multiple islands. Takenaka et al. [17] identified genetic differentiation among nine major populations according to island, based on a combination of partial mtDNA and nuDNA regions (COI, 16S rRNA, ITS, and histone H3), suggesting that straits between islands act as major genetic barriers for *G*. *dehaani*. In contrast, our results highlight the importance of geological events, including volcanic activity and climate change, in shaping the genetic structure of this species group.

For example, the co-occurrence of three populations (nKC, cK, and sKK) on Kyushu Island cannot be explained solely by the geographical barrier posed by the strait, as these populations are separated along a North–South axis with distinct geographical boundaries. ABC analysis suggested that they diverged nearly simultaneously from a common ancestor, implying a large-scale or continuous event disrupted gene flow across Kyushu. Moreover, the identified boundaries appear to be influenced by sea level fluctuations and volcanic activity. For example, the nKC population spans the Kanmon Strait into central Chugoku and northern Kyushu. Despite its unclear boundary with the HO population, it aligns with the distribution boundaries of other species, such as calopterygid damselflies and small salamanders [29, 30], hinting at historical events in the Chugoku region that impeded gene flow. Major rivers, such as Gohnokawa and Ota, which flow across the Chugoku Mountains to the Sea of Japan, may have once served as barriers connected to the Seto Inland Sea [31]. In contrast, the southern boundary between the nKC and cK lacks a singular geographical barrier, suggesting that multiple factors influenced their genetic divergence. Submerged plains, such as Tsukushi, during past interglacial periods and the adjacent Sefuri Mountains may have partially restricted gene flow in northern freshwater organisms [27, 32]. Additionally, volcanic activity between 5–6 MYA and 0.5–2.0 MYA along the Beppu–Shimabara Graben, which runs East–West through central Kyushu [33], likely shaped local barrier and limited gene flow. In fact, the latter period activity may have influenced the amphibian distribution [34]. Such disturbances promoted local isolation and genetic differentiation. Similarly, the cK–sKK boundary appears to be influenced by volcanic activity, as noted by Suzuki & Tsuda [23].

The HO, SHI, and sKK populations exhibited a unique distribution pattern, with the latter two clearly displaying enclave distributions. Two scenarios could explain such distribution: a stable contact zone scenario, in which a population persists as a remnant patch of a historically widespread distribution, or a moving contact zone scenario, in which a population migrates into new areas [35]. The SHI population likely aligns with the first scenario for several reasons: (1) geographic and ecological barriers make recent colonization unlikely, (2) exhibits low genetic diversity (π = 0.057), suggesting a historical bottleneck, and (3) evidence of rapid expansion in the HO population by low genetic diversity in SNPs (π = 0.059) and neutrality tests, including a negative Tajima’s *D* (−1.69) and Fu’s *Fs* (−23.7), imply competitive displacement. Admixture analysis further supports this hypothesis, revealing the SHI population share ancestry with the HO population in both the Kii Peninsula and Shikoku Island (Fig. 3). These findings suggest that the SHI population once extended across these regions but got fragmented as the rapidly expanding HO population isolated SHI individuals into enclaves. In contrast, presence of the HO population in northwestern Shikoku represents a moving contact zone scenario driven by its rapid expansion. Notably, the HO population exhibits the widest distribution among all, extending as far as Hokkaido and crossing the Blakiston Line, a key faunal boundary. Our results revealed no significant genetic differences in mtDNA and nuDNA between the Hokkaido specimens and other HO specimens, indicating either recent expansion or human-mediated introduction, as proposed by Sugime et al. [11]. These findings highlight the dynamic role of the HO population in shaping *G*. *dehaani* population distribution through rapid expansion, and its influence on the enclave distribution of other populations.

Similarly, the sKK population likely aligns with the moving contact zone scenario. This population, found in the southern Kyushu, nearby islands, and along the coastal areas from the Izu Peninsula to the Boso Peninsula (southern Kanto), exhibits a distinctive distribution pattern. Takenaka et al. [17] hypothesized that the population in southern Kyushu and southern Kanto (clades 3e, 3f, 3g, and 3h in this study) expanded via oceanic dispersal when the Izu Peninsula was an island. Although our SNP-based results appear to support this hypothesis, the limited genetic diversity data of the sKK population prevent confirmation at this stage. Although the admixture analysis grouped specimens from southern Kyushu and southern Kanto into a single sKK population, the COI-based neighbor network (Fig. 2) revealed genetic differences between these clades (clades 3e, 3f, 3g, and 3h). This indicates the possibility of a relictual distribution for the southern Kanto subgroup. Future studies employing high-resolution methods, such as whole-genome sequencing, are necessary to clarify their intrapopulation genetic structure and provide deeper insights into the unique enclaved distribution of the sKK population.

### Relationship between body color trait and genetic population structure

This study did not find a complete correspondence between body color types (DA, RE, BL, OC, and amakusa) and populations using COI and SNP analyses. However, we observed a biased distribution pattern along Pacific coast areas for the BL and RE types. Although the BL type emerged twice independently in the evolutionary history of the *G. dehaani* species complex (Fig. 4, sKK and SHI populations), such a biased distribution of the BL type may be due to local climate adaptation. Pale carapace coloration in crustaceans may play a thermoregulatory role [36]; furthermore, the range of the BL type overlaps with warm Pacific coastal areas influenced by the Kuroshio Current. A similar pattern was observed for the RE type, with individuals concentrated along the Pacific coast, compared to the DA types (Fig. 3). Although COI and SNP data revealed no genetic differences between DA and RE types within the same populations, geographical distribution of prey having astaxanthin precursors, such as β-carotene and zeaxanthin [23, 37], may influence this bias. Alternatively, an undetected genetic structure within the population may be another contributor. Despite the unclear mechanism behind the biased distribution patterns of the BL and RE types, our admixture analysis and body color distribution mapping highlight the potential of the *G. dehaani* species complex to serve as a model for studying local adaptation to climate and environment.

Despite the incomplete correspondence between body color traits and populations, combining body color with collection locality data could help identify the population affinities of collected specimens. For example, the sKK population, exclusively comprising BL individuals, spans southern Kyushu and southern Kanto, indicating that body color and collection locality can together distinguish populations. Similarly, the HO and nKC populations, predominantly of DA or RE type, can often be identified this way, despite rare BL-type individuals (Table S1, No. 249 in the HO population area and No. 313 in the nKC area). These rare cases likely represent an individual-level color variant, considering the limited collection sites and specimen numbers, similar to findings for *Ryukyum yaeyamense* [38].

In contrast, the SHI and cK populations exhibited greater intraspecific variation in body coloration (Figs. 1, 3). Although the SHI population predominantly comprises BL individuals, it also includes DA/RE types. Mismatched specimens sharing multiple common ancestors (Fig. 3) suggest that the genetic introgression through secondary contact contributes to the discordance in body color within the population. The cK population also displayed various coloration types, including DA, RE, OC, and even *G*. *amakusa* types, despite consistent genetic proportions (Fig. 3). Such variations in SHI and cK populations imply historical or ongoing gene flow and highlight the need for reevaluating the genetic and morphological boundaries in the *G. dehaani* species complex, including *G*. *amakusa*. Molecular methods, especially genome-wide analyses, are essential for resolving the genetic structure and identifying individuals in SHI and cK effectively in future research.

### Discordance between mtDNA and nuDNA sequences

Our molecular analysis highlights the need for caution when interpreting previous studies relying on the combinations of partial mtDNA and nuDNA sequences. While our COI-based neighbor-net diagram aligns with earlier low-resolution studies using isozymes or partial sequences [14, 15, 17, 27], the population structure revealed by our ADMIXTURE analysis using SNPs significantly differs. This discordance emphasizes the importance of genome-wide analyses alongside partial sequence analyses for accurately elucidating the genetic phylogeny and population structure of the *G. dehaani* species complex.

Discordances between mtDNA and nuDNA are common in various taxa and often attributed to adaptive mtDNA introgression, demographic disparities, or sex-related asymmetries [20]. In our study, the *G. dehaani* species complex was divided into three clades based on mtDNA, whereas nuDNA identified five populations. In areas of population overlap, individuals exhibited genetic elements from both populations (e.g., nKC and HO populations; Fig. 3), suggesting that contact between the two promotes gene flow and contributes to the discordance between mtDNA and nuDNA results.

mtDNA populations are widely distributed as enclaves (e.g., pop2: Kyushu and Shikoku, pop3d: northeastern Shikoku and Kyushu), whereas nuDNA populations are divided into SHI, nKC, cK, and sKK, with clear regional divisions. For the many taxa where mtDNA and nuDNA do not match, mtDNA is less structured than nuDNA (Toews & Brelsford, 2012); a similar result was obtained for the *G. dehaani* species complex. The complex mtDNA distribution pattern is probably due to repeated isolation and merge of populations due to past geographical changes. Over time, nuDNA may have become uniform within populations while mtDNA variation due to hybridization or ecological differences, such as sex-biased mobility, was maintained.

Our research indicates that gene flow may still be ongoing or might have historically occurred but has since ceased. This study did not analyze the gene flow dynamics in detail, as the samples were selected to investigate the overall genetic characteristics of *G. dehaani* populations. Future studies focusing on genetic boundary regions are needed to clarify these patterns.

## Conclusion

Our study reveals the complex genetic population structure of the Japanese freshwater crab *G. dehaani* species complex, resulting from multiple geological events, including volcanic activity and sea level fluctuations, in the Japanese Islands. Admixture analysis identified five populations with distinct geographic boundaries. On Kyushu Island, three distinct populations (nKC, cK, and sKK) separated along a North–South gradient were likely influenced by large-scale disturbances, such as volcanic activity. The recent rapid expansion of the HO population, indicated by low genetic diversity, has contributed to the unique enclave distribution of the SHI population. Further, regional body color variations partially correlate with SNP clades, suggesting the potential of combining collection locality and body color for population identification. However, further molecular and morphological approaches are needed for precise population-identification from specimens. Contrasting patterns between mtDNA and nuDNA highlight historical gene flow and adaptive introgression, underscoring the need for comprehensive genomic analysis. In addition, the discordance between nuclear SNPs and mtDNA classifications warrants caution when interpreting earlier phylogenetic studies based on combined mtDNA and nuDNA data. These findings provide new insights into the evolutionary history and genetic diversity of the *G. dehaani* species complex, providing a basis for future research on its taxonomy, phylogeny, and population dynamics.

## Materials and methods

### Sample collection and laboratory procedures

From 2019 to 2023, 1037 *Geothelphusa* specimens were collected from 299 sites, spanning from Hokkaido to the central Ryukyu Islands (Table S7). In this study, *G*. *dehaani*, *G*. *mutsu*, and *G*. *amakusa* were all considered as *G. dehaani* species complex without distinction due to ambiguous species boundaries and limited distribution knowledge. Almost all specimens were transported to the laboratory either alive or anesthetized, and photographs were captured to document live body coloration. After color correction using color bars, specimens were categorized into four types based on Chokki [21, 22] and Naruse and Ng [9]: BL (grayish blue or bluish green carapace, occasionally including dark reddish purple, with light grayish or light yellow thoracic legs), DA (dark purplish brown carapace and thoracic legs), RE (dark brown carapace, sometimes with orange or brown color on the posterolateral portion, and reddish orange thoracic legs), and “amakusa” (a sky blue carapace with whitish, brownish, or reddish thoracic legs). Specimens with unclassifiable color patterns were labeled as OC (other color). Specimens were fixed in 80% ethanol, and part of a leg was transferred to 99% ethanol for DNA analysis. Some specimens were directly fixed in 99.5% ethanol in the field. In addition, out-group specimens, including *G. exigua* (n = 2, collected at southern Kyushu), *G*. *marmorata* (n = 2, Yakushima Island), *G*. *koshikiensis* (n = 6, Koshiki Islands), and *G*. *sakamotoanus* (n = 2, Okinawa Island and Tokunoshima Island) were treated similar to *G. dehaani* samples. The identification of out-group specimens was based on morphological characteristics, such as body color and male first pleopod shapes [10]. All fixed specimens were registered and are preserved at the Wakayama Museum of Nature (WMNH). Our research protocols complied with the current laws of Japan, including the animal welfare ones, and followed the guidelines for protecting and promoting decapod crustacean welfare in research, by the Insect Welfare Research Society [39].

### DNA extraction, sequencing, and alignment

**mtDNA.** A total of 509 specimens from 219 localities were selected for mtDNA analysis of the COI region, covering the distribution ranges of the *G. dehaani* species complex (Table S1). Genomic DNA was extracted from the leg muscles of each specimen using a DNeasy Blood and Tissue Kit (Qiagen, Hilden, Germany). Partial sequences of the mtDNA COI-encoding region were amplified by PCR using the universal primers jgLCO1490 and jgHCO2198 [40] with TaKaRa Ex Taq (TaKaRa, Shiga, Japan). The PCR protocol involved mixing 10 µL of Premix TaqTM (Takara Bio Inc., Tokyo, Japan), 1.2 µL of each primer (10 µM), 6.6 µL of Milli-Q water, and 1 µL of template DNA. PCR was performed using Applied Biosystems VeritiPro (Thermo Fisher Scientific K.K., Massachusetts, USA) under the following conditions: 94°C for 1 min, followed by 35 cycles of 94°C for 40 s, 51°C for 40 s, and 72°C for 1 min, and final elongation at 72°C for 7 min. A 3-µL sample of the PCR product was mixed with 1 µL of Midori Green Direct DNA Stain (NIPPON Genetics, Tokyo, Japan) and electrophoresed on a 2% agarose gel at 100 V for 15 min. Then, the gel was visualized under UV light to confirm DNA amplification. Amplification products were purified by mixing 17 µL of PCR products with 1 µL of ExoSAP-IT solution (Thermo Fisher Scientific K.K., Massachusetts, USA) and incubating at 37°C for 15 min and 80°C for 15 min. The purified PCR products were then sent to Macro-gen Japan (Tokyo, Japan) for sequencing with the same primers used for PCR amplification. The resulting sequences were aligned using the MEGA 11 and ClustalW software [41]. The mtDNA sequences were registered in the DNA Data Bank of Japan (DDBJ) (accession numbers: LC864606–LC865102), and the haplotypes were determined (Table S2).

**SNPs**. Genome-wide SNPs were analyzed using multiplexed inter-simple sequence repeat genotyping by sequencing (MIG-seq) [42] for 154 individuals from 92 sites in the *G. dehaani* species complex utilizing the Illumina Hiseq system (Illumina) (Table S1). To ensure high-quality sequence data, reads 1 and 2, adapter regions, and low-quality reads were removed using fastp ver. 3 [43]. The quality threshold was set to 30, with a required length of 134, trimming front 1 and front 2 by 14, cutting front/tail set to 20. The law data obtained by MIG-seq were registered at DDBJ (accession numbers: DRR641568–DRR641721).

### Population genetic analysis mtDNA analysis

In addition to the samples used in this study, Genbank samples from *G*. *dehaani* from Hachijyo Island (LC743147, LC743148), *G*. *dehaani* from Kuchinoshima Island (LC743174) [17], *G*. *dehaani* from Naju, South Korea (MG674171) [44], and *G. sakamotoanus* from Takara Island (LC743301, LC743302) [17] were included in the analysis. To assess genetic diversity and neutrality, haplotype diversity (*h*), nucleotide diversity (π) [45], Tajima’s *D* test [46], and Fu’s *Fs* test [47] were calculated for each population using Arlequin version 3.5.2.2 [48]. Genetic differentiation among populations was examined using the pairwise fixation index (*F*_ST_) and *F*_ST_ *p*-value [49]. A network diagram was created using the neighbor-net method [50] with SplitsTree4 [51].

### SNP analysis

Quality-filtered sequence data were used to identify SNPs using Stacks 2.41 [52]. The Stacks pipeline denovo_map.pl was implemented with default values, except where indicated. In ustacks, the minimum coverage depth (m) was set to 5 and the maximum distance between stacks (M) to 2. For population analysis, the min-samples-overall (R) was set to 0.5, the minimum minor allele count (min-mac) to 2, and the maximum observed heterozygosity (max-obs-het) to 0.75; only the first SNP per locus was used. We calculated the *Ho*, *He* [45], π, *F*_IS_ [53], *F*_ST_, and *F*′_ST_ [54] using the fstats module in the populations program in Stacks 2.41.

Genetic structure analysis was performed with ADMIXTURE version 1.3.0 [55], using C set to 0.0001 and S set to time. Output data were visualized in Excel. All individuals included in the SNP analysis (set 0) were analyzed by dividing the poplar into three mtDNA clades. Based on the genetic structure analysis of all individuals, five populations were identified, and ADMIXTURE analysis was performed on six adjacent sets to assess hybridization (Table S5). Set 1 included populations in the Kanto region (17 individuals from Ibaraki, Tochigi, Gunma, Saitama, Tokyo, Kanagawa, and Shizuoka prefectures). Set 2 covered the Kinki region (16 individuals from Shiga, Mie, Nara, and Wakayama prefectures) and set 3 the Chugoku region (16 individuals from Kyoto, Hyogo, Tottori, Okayama, Shimane, Hiroshima, and Yamaguchi Prefectures), while set 4 included populations in the Shikoku region (21 individuals from Awaji Island-Hyogo Prefecture, Kagawa, Tokushima, Kochi, and Ehime Prefectures), set 5 represented central Kyushu (31 individuals from Fukuoka, Saga, Oita, Nagasaki, and Kumamoto Prefectures), and set 6 included populations in southern Kyushu (39 individuals from Saga, Nagasaki, Kumamoto, Miyazaki, and Kagoshima Prefectures). These poplars were analyzed to evaluate the genetic structure derived from SNP results.

### Estimation of the population demography history

To investigate the history of genetic diversification, we conducted a coalescent analysis using ABC with DIYABC Random Forest version 2.1 [56]. Based on the results of our STRUCTURE analysis (see results), we classified the populations into five groups: pop1 (Pacific coast from Boso Peninsula to Izu Peninsula + southern Kyushu and surrounding islands, hereafter referred to as the sKK population), pop2 (southern Shikoku + part of Kii Peninsula, SHI population), pop3 (central Kyushu, cK population), pop4 (northern Kyushu + Chugoku region of Honshu, nKC population), and pop5 (Hokkaido + Honshu from Tohoku to Chugoku + part of Shikoku, HO population). As ABC frameworks require populations without continuous gene flow, we focused the analysis on 118 specimens likely descending from a single ancestral population. We used the 313 SNP loci for this analysis, because we excluded SNPs not present in ≥1 sample. We used Hudson’s algorithm [57] to calculate the minor allele frequency. Assuming a generation time of four years for *G. dehaani* [58] and an Early Pleistocene age (2 Mya) for the ancestral population’s initial colony formation [59], we set the effective population sizes and generation parameters a priori at 100,000–500,000 (Table S7). All prior values were drawn from uniform distributions, under the condition t1 < t2 < t3 < t4.

For scenario selection, the DIYABC assigns a vote for each scenario, reflecting how often it was chosen across a forest of n trees. The scenario with the highest vote count is considered the best fit for the dataset, alongside an estimate of the posterior probability for that scenario. A training set comprising 64,000 simulations and 500 trees was used to identify the most supported model scenario. Prior to that, we performed pre-scenario checks using PCA to detect potential model misspecification by comparing the prior and posterior parameter distributions.

In this study, we evaluated a total of 32 scenarios across eight groups (Figure S3): 1, scenarios based on body color (i.e., BL vs. DA, scenarios 1–3); 2, scenarios based on geographic separation areas at first (i.e., Honshu vs. Kyushu, scenarios 4–7); 3–7, scenarios where each population diverged sequentially (i.e., from sKK to HO, scenarios 8–11, 12–15, 16–19, 20–23, and 24–28, respectively); and 8, scenarios involving nearly simultaneous multiple divergences (scenarios 29–32).

### Data availability

Raw data about collection localities of the *G*. *dehaani* species complex are available in supplementary information. The detailed information, however, was excluded from the viewpoint of habitat conservation.

## Acknowledgments

We are sincerely grateful to the following people helped sample collections: R. Asada, K. Eto, S. Fuji, K. Fukutani, I. Fukuyama, R. Fukuyama, K. Hamanaka, S. Hara, R. Hayashi, Y. Hibino, K. Hirashima, K. Hirose, M. Iida, T. Iwasaki, A. Kanamori, S. Kanamori, K. Kawamura, A. Kimoto, K. Kirihara, M. Kirihara, O. Kishida, T. Kita, H. Kobayashi, Y. Kojima, R. Kudo, S. Matsuno, N. Miyajima, Y. Mizoguchi, Y. Murase, K. Natsukawa, K. Niwa, Y. Obae, K. Okada, K. Okazaki, T. Sato, Y. Shen, C. Shinomiya, R. Sugime, A. Tagawa, Y. Tomimori, T. Toyama, K. Toyota, U. Yamakawa, Y. Yamamoto, H. Yanagi, and R. Yoshida. In addition to the samples used in this study, we also received samples from many other people. Members of the Nishikawa Laboratory at Kyoto University provided comments on the analysis. The sample of the Danjo Islands provided by Dr. K. Eto was investigated under the following permit: “2-ju Bun-Cho No. 4-173” (Name of change permit: Kitakyushu Museum of Natural History and Human History (Director) / Person in charge of change: Eto). The research on Yakushima Island was conducted with permission from the Kyushu Regional Forest Agency for K.T.

## Author contributions

T. K. and K.T. share to the first and corresponding authors, with equal to contribution. T.K. and K.T. designed the study, analyzed the data, visualized the results, and wrote the original manuscript. T.K., K.T., and K.N. collected the specimens. K.T., K.G., M.N. and A.I. conducted the molecular analyses. All authors reviewed the draft and approved the final manuscript.

