## Supplementary material for "Genetic population structure of Japanese freshwater crab, *Geothelphusa dehaani* species complex (Decapoda: Potamidae) using genome-wide SNP": Fig. S1

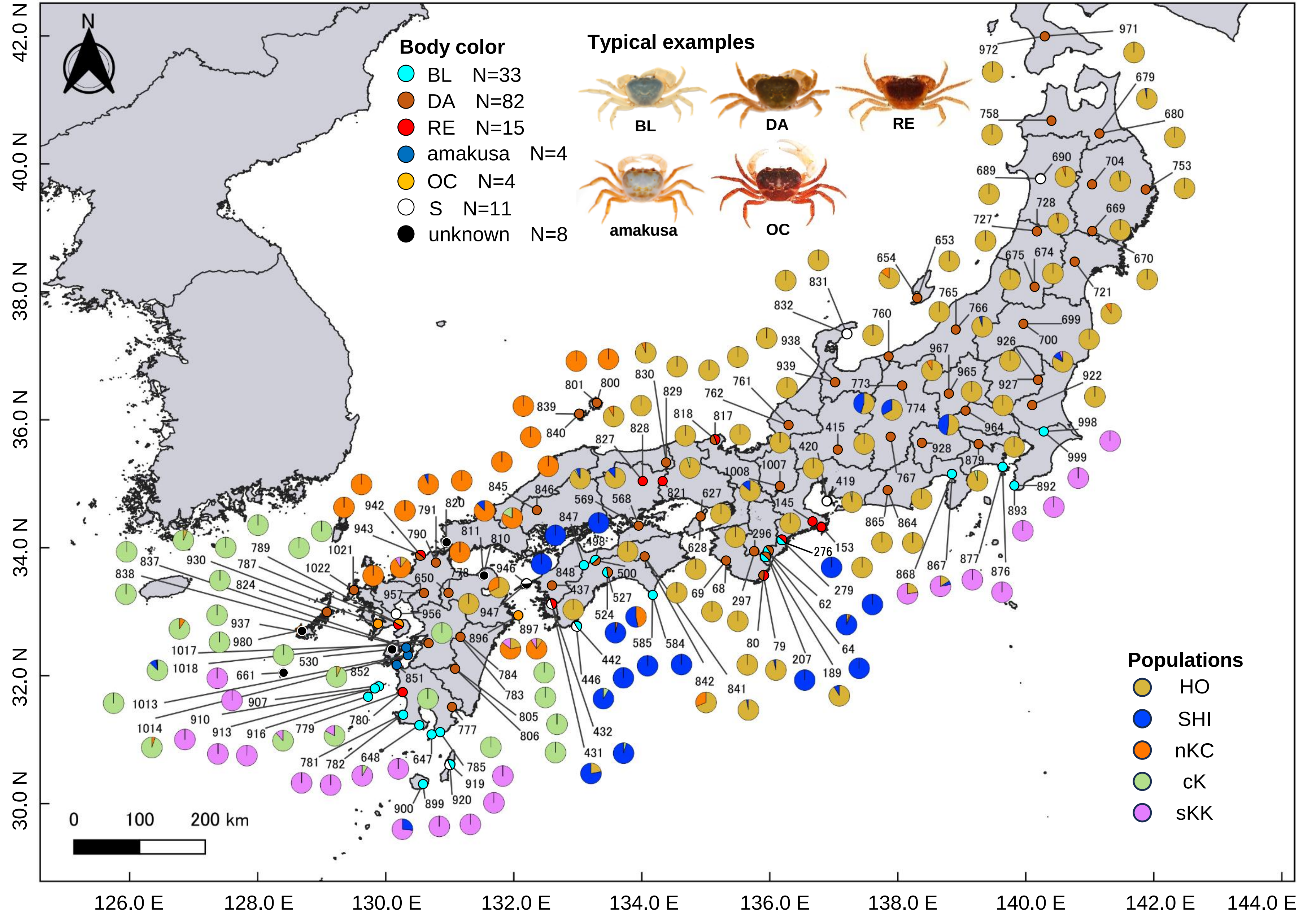


Figure S1. Map showing detailed geographical distributions of color types with sample numbers, and the results of the admixture analysis for each location (set 0). Large pie chart, genetic elements of each individual; small pie chart, color types of individuals at each location used in this analysis. Color types are indicated as in Figure 1. The colors in the pie charts corresponding to each group and genetic elements are as follows: HO, brown; SHI, blue; nKC, orange; cK, green; and sKK, pink. The sample numbers refer to Table S1, Figs. 1 and 3.
