## Supplementary material for "Genetic population structure of Japanese freshwater crab, *Geothelphusa dehaani* species complex (Decapoda: Potamidae) using genome-wide SNP": Fig. S2

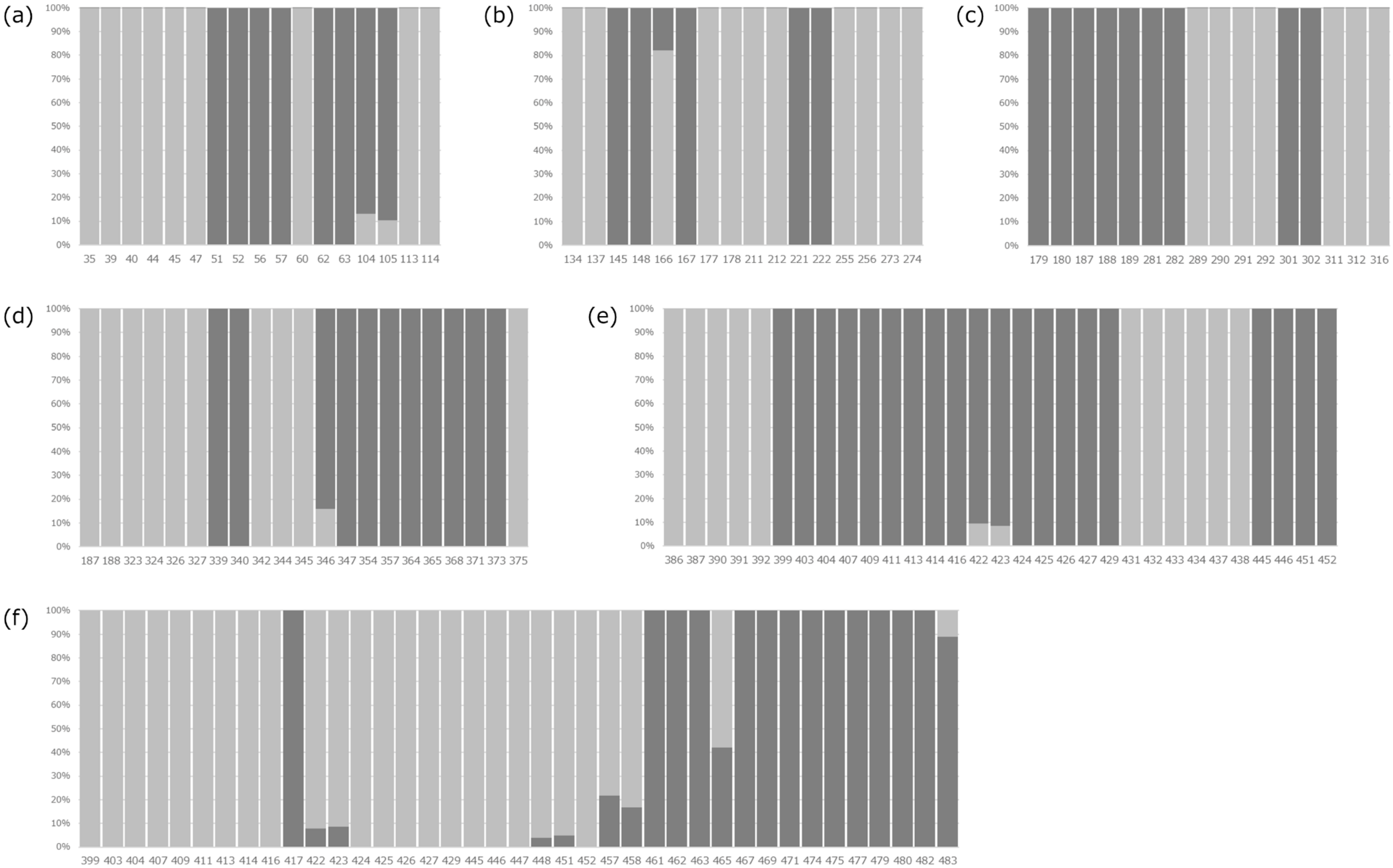


Figure S2. Admixture analysis using samples from the two populations at each group boundary. For sample numbers, refer to Table S1. (a) Set 1: Kanto region (17 individuals from Ibaraki, Tochigi, Gunma, Saitama, Tokyo, Kanagawa, and Shizuoka prefectures), (b) Set 2: Kinki region (16 individuals from Shiga, Mie, Nara, and Wakayama prefectures), (c) Set 3: Chugoku region (16 individuals from Kyoto, Hyogo, Tottori, Okayama, Shimane, Hiroshima, and Yamaguchi Prefectures), (d) Set 4: Shikoku region (21 individuals from Awaji Island-Hyogo Prefecture, Kagawa, Tokushima, Kochi, and Ehime Prefectures), (e) Set 5: central Kyushu (31 individuals from Fukuoka, Saga, Oita, Nagasaki, and Kumamoto Prefectures), and (f) Set 6: southern Kyushu (39 individuals from Saga, Nagasaki, Kumamoto, Miyazaki, and Kagoshima Prefectures).
