## Supplementary material for "Genetic population structure of Japanese freshwater crab, *Geothelphusa dehaani* species complex (Decapoda: Potamidae) using genome-wide SNP": Fig. S3

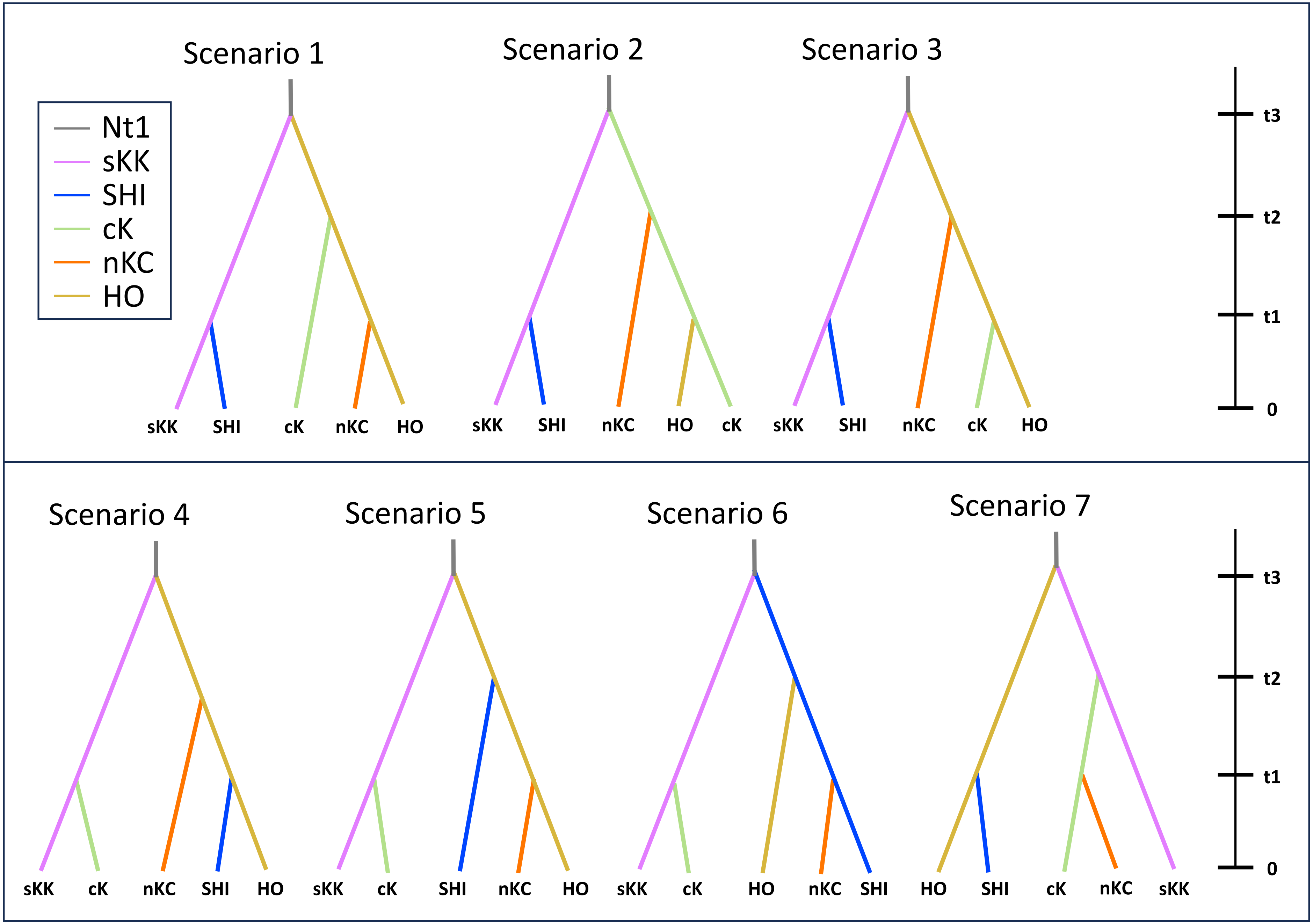


Figure S3. Tested demographic scenarios in the DIYABC-RF analysis for the five *Geothelphusa dehaani* population datasets. See detailed parameters in Table S7.

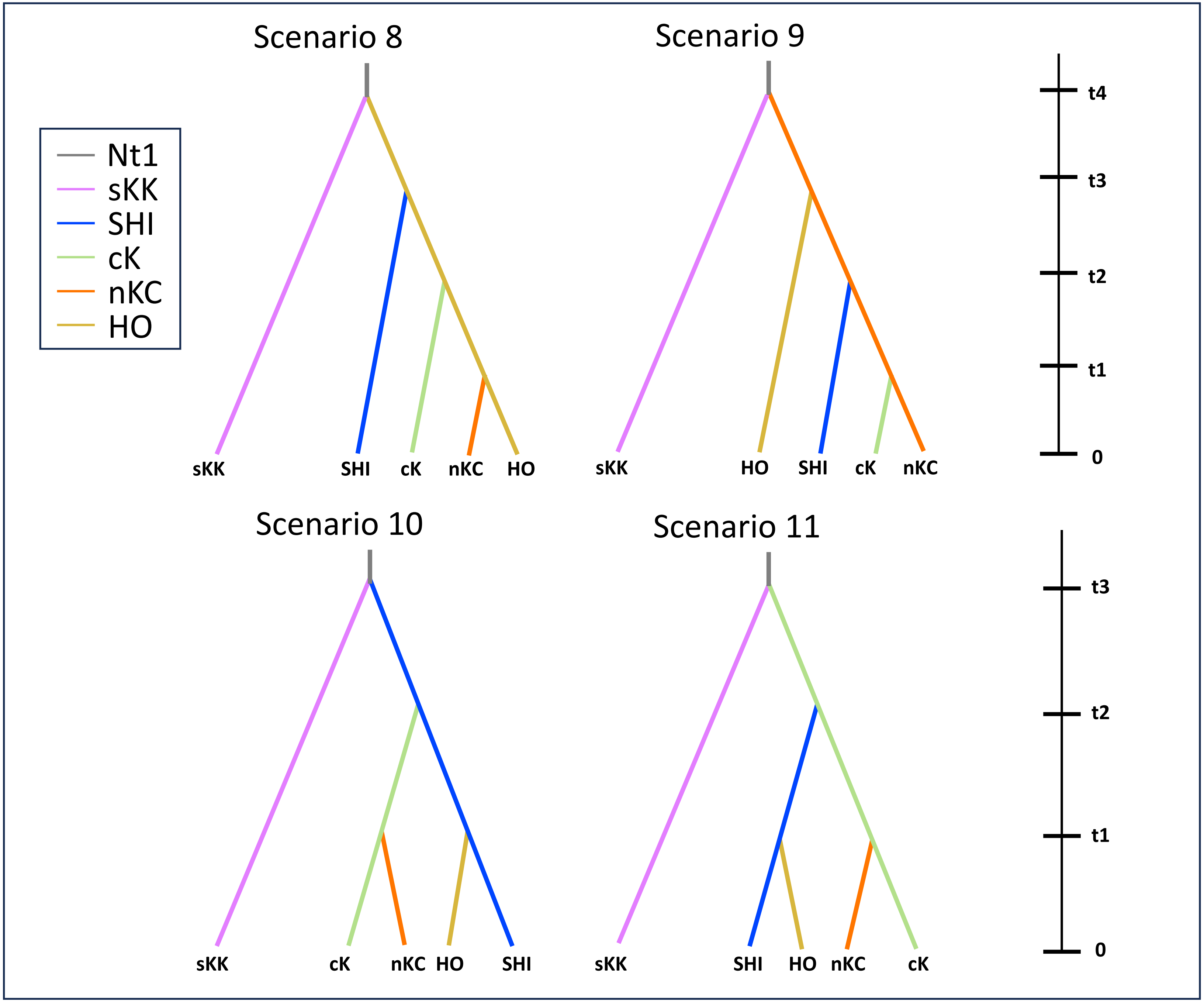


Figure S3 Continued.

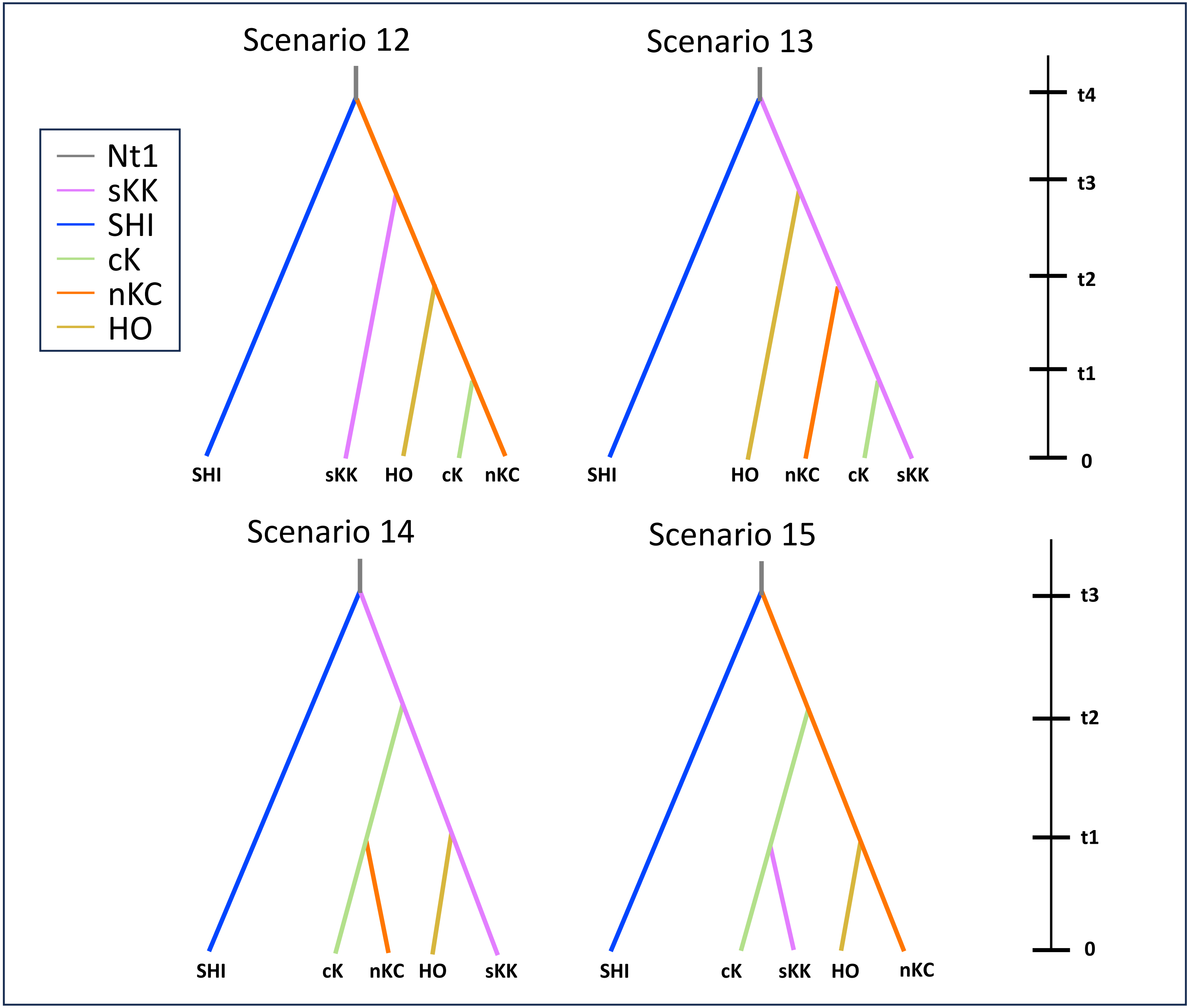


Figure S3 Continued.

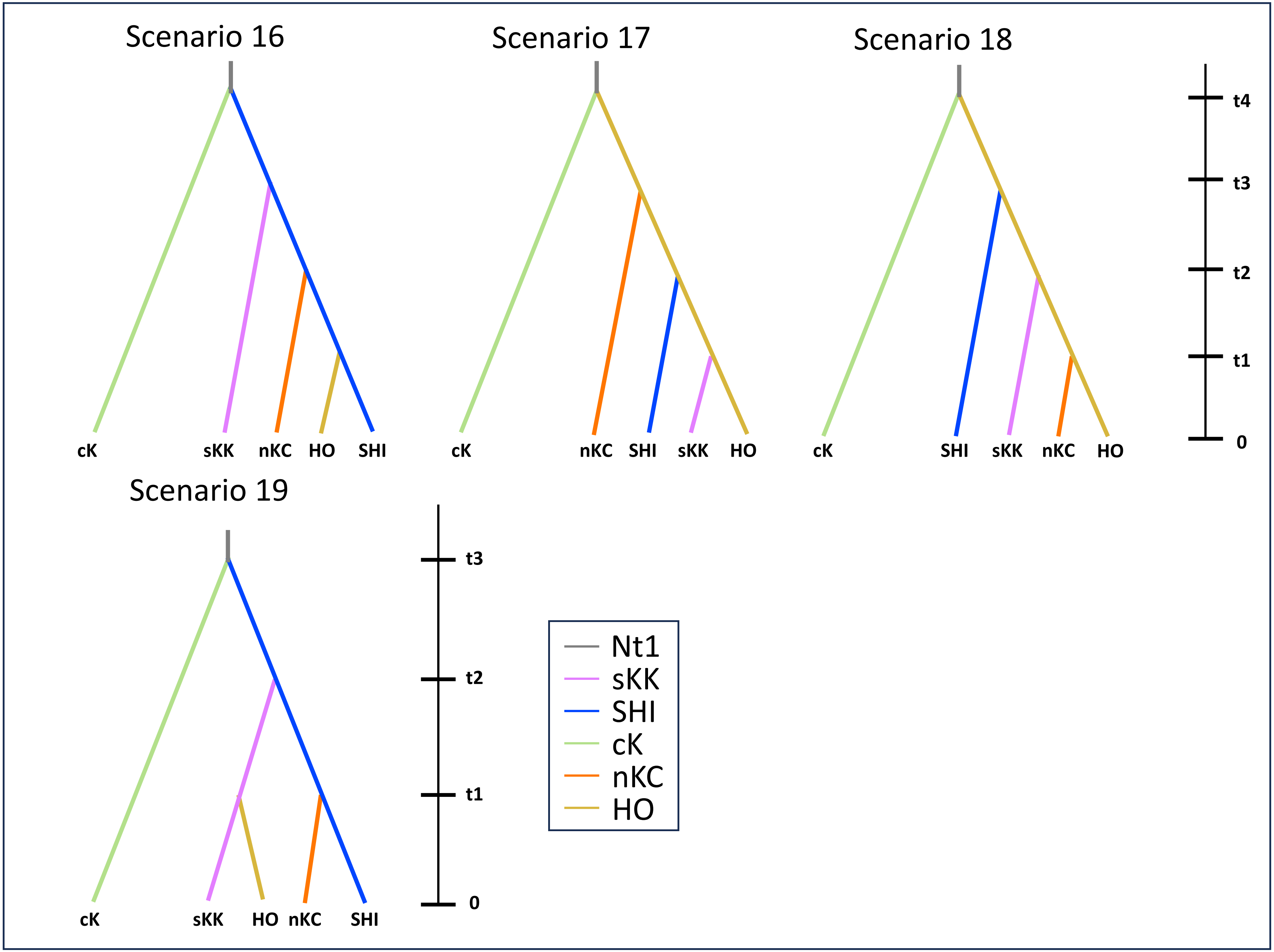


Figure S3 Continued.

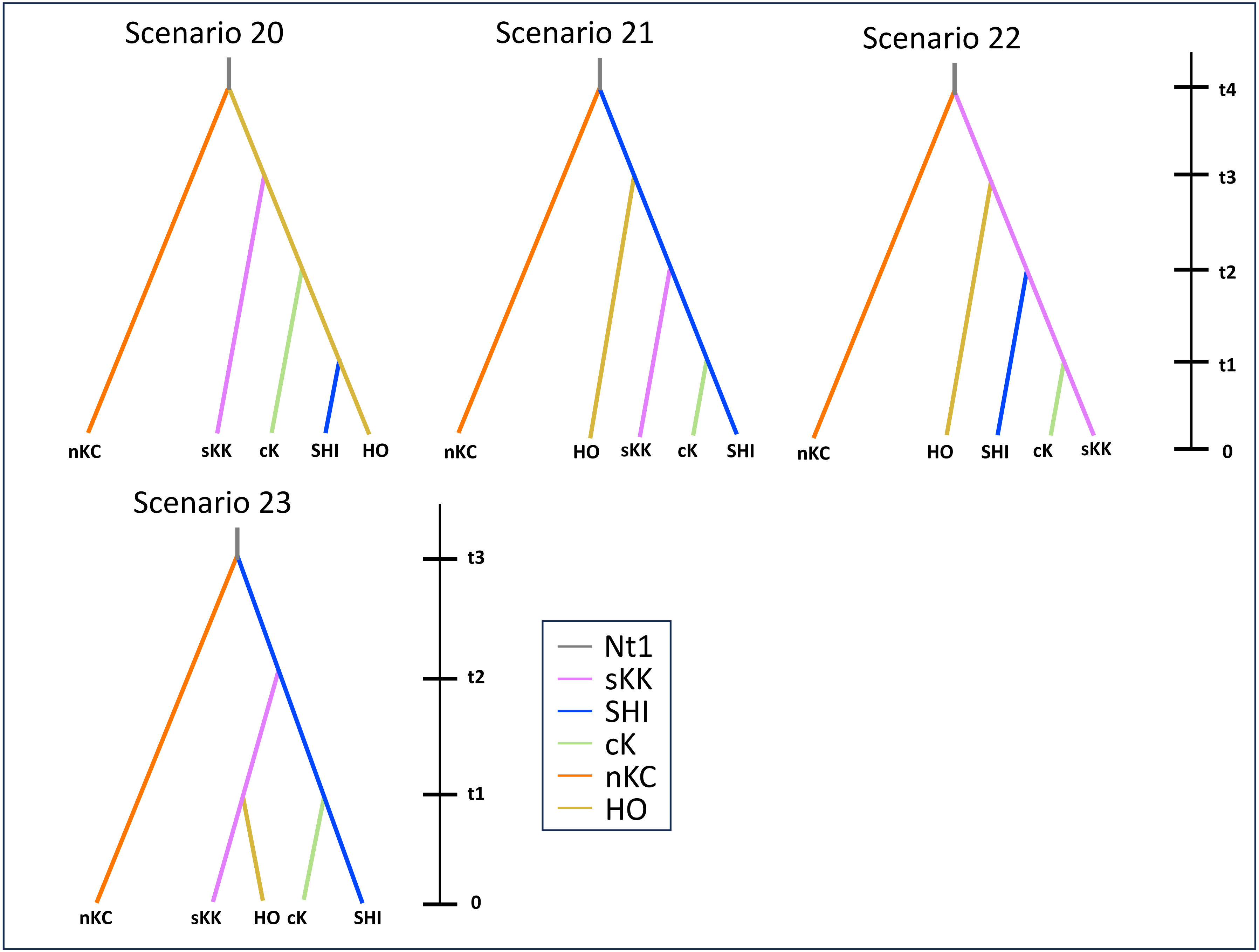


Figure S3 Continued.

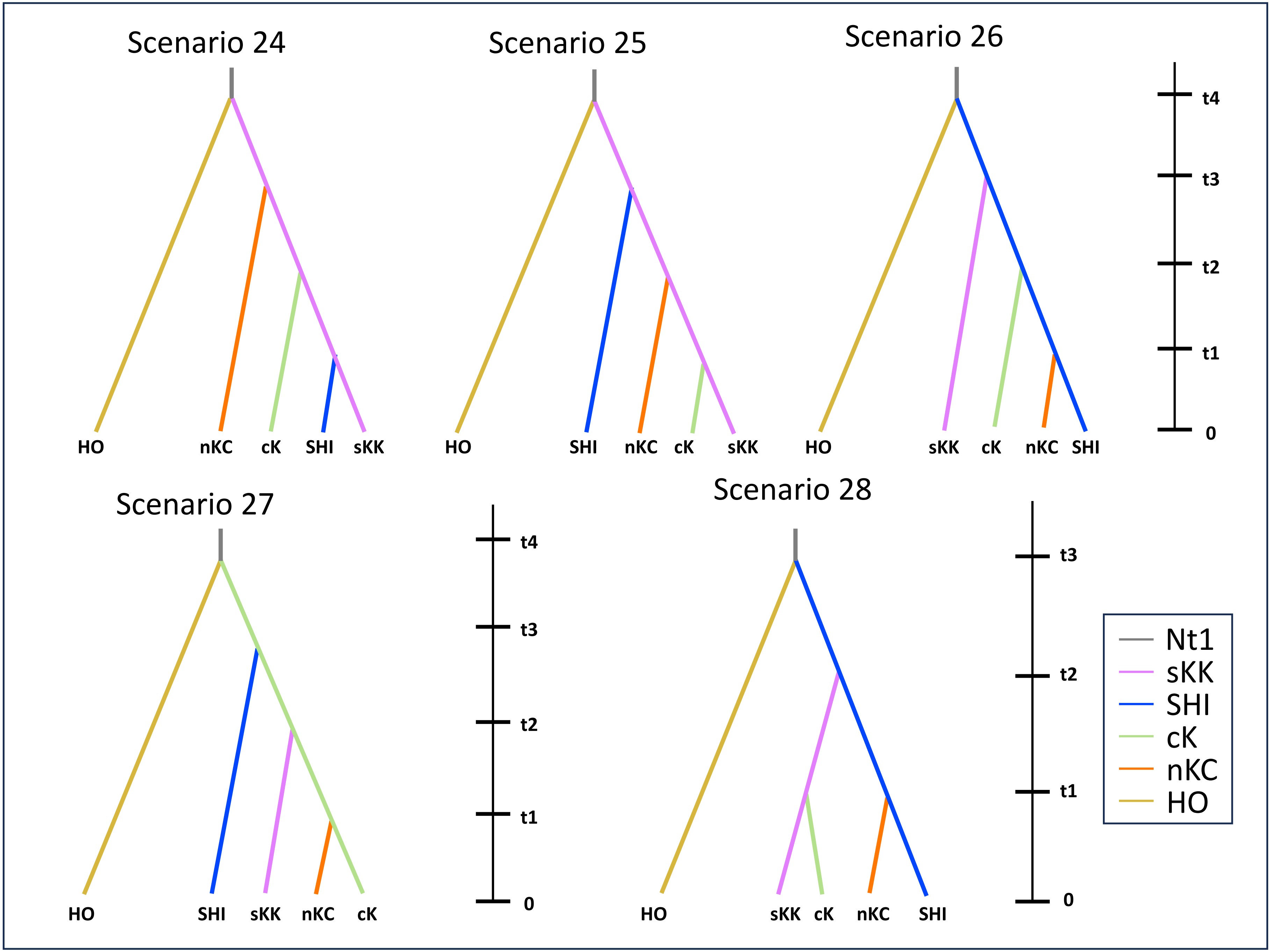


Figure S3 Continued.

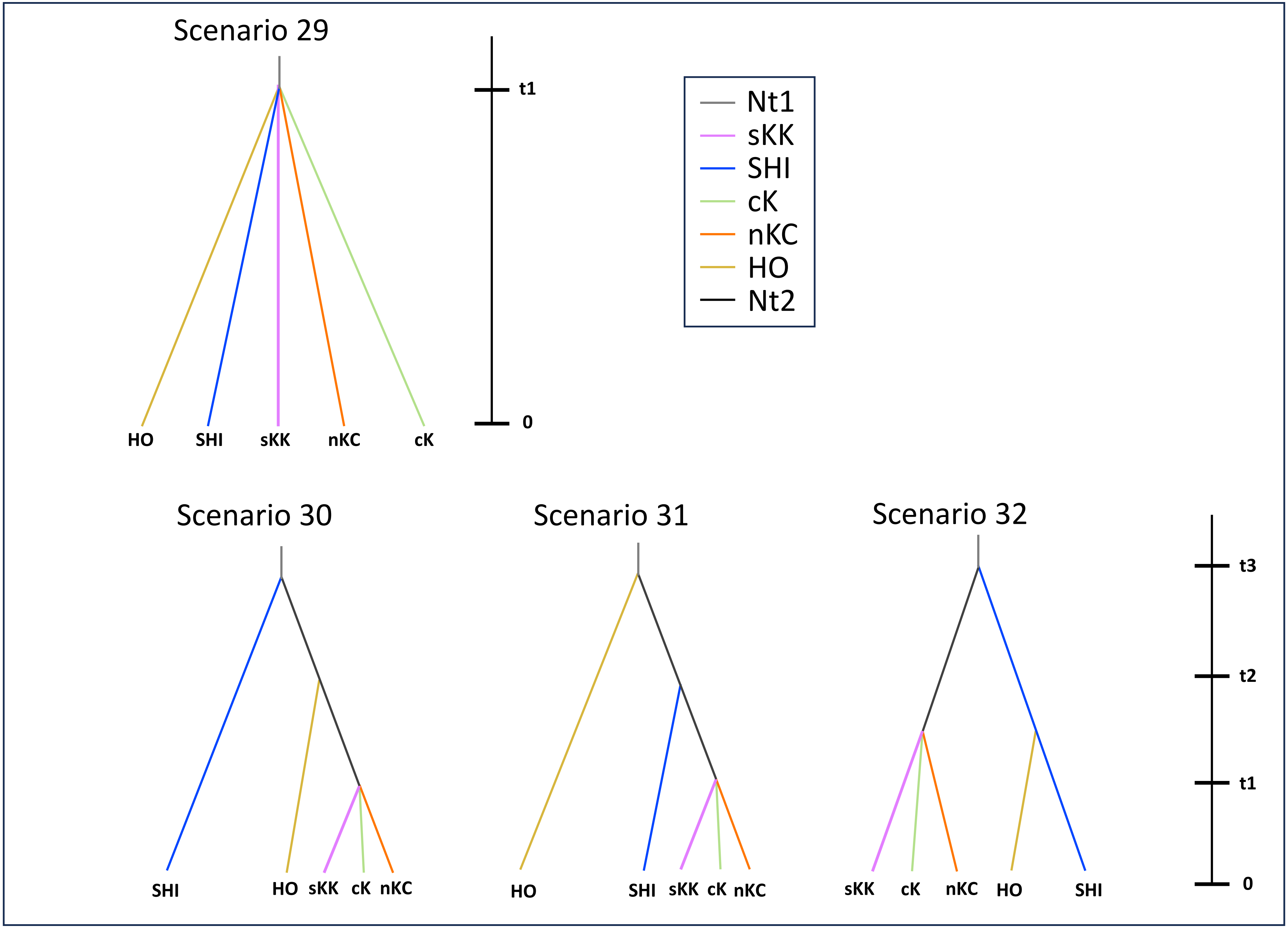


Figure S3 Continued.
