## Supplementary material for "Genetic population structure of Japanese freshwater crab, *Geothelphusa dehaani* species complex (Decapoda: Potamidae) using genome-wide SNP": Table S1

Table S1. List of sampling and sequencing information.

| No. | WMNH-KT- | Species Name | Prefecture | City | Other | Color Type | mtDNA Accesion number | mtDNA Group number | SNPs Accesion number |
|---|---|---|---|---|---|---|---|---|---|
| 1 | 968 | *G. dehaani* | Hokkaido | - | Oshima Peninsula | DA | LC864606 | 3a | - |
| 2 | 969 | *G. dehaani* | Hokkaido | - | Oshima Peninsula | DA | LC864607 | 3a | - |
| 3 | 970 | *G. dehaani* | Hokkaido | - | Oshima Peninsula | DA | LC864608 | 3a | - |
| 4 | 971 | *G. dehaani* | Hokkaido | - | Oshima Peninsula | DA | - | - | DRR641709 |
| 5 | 972 | *G. dehaani* | Hokkaido | - | Oshima Peninsula | DA | LC864609 | 3a | DRR641710 |
| 6 | 757 | *G. dehaani* | Aomori | Hirosaki | Onizawa | DA | LC864610 | 3a | - |
| 7 | 758 | *G. dehaani* | Aomori | Hirosaki | Onizawa | DA | LC864611 | 3a | DRR641622 |
| 8 | 679 | *G. dehaani* | Aomori | Shingo | Herai | DA | LC864612 | 3a | DRR641611 |
| 9 | 680 | *G. dehaani* | Aomori | Shingo | Herai | DA | LC864613 | 3a | DRR641612 |
| 10 | 684 | *G. dehaani* | Iwate | Kuji | Natsui | DA | LC864614 | 3a | - |
| 11 | 685 | *G. dehaani* | Iwate | Kuji | Natsui | DA | LC864615 | 3a | - |
| 12 | 753 | *G. dehaani* | Iwate | Miyako | Nagasawa | DA | LC864616 | 3a | DRR641621 |
| 13 | 754 | *G. dehaani* | Iwate | Miyako | Nagasawa | S | LC864617 | 3a | - |
| 14 | 704 | *G. dehaani* | Iwate | Morioka | Tsunagikitakubo | DA | LC864618 | 3a | DRR641617 |
| 15 | 705 | *G. dehaani* | Iwate | Morioka | Tsunagikitakubo | DA | LC864619 | 3a | - |
| 16 | 706 | *G. dehaani* | Iwate | Morioka | Tsunagikitakubo | DA | LC864620 | 3a | - |
| 17 | 707 | *G. dehaani* | Iwate | Morioka | Tsunagikitakubo | DA | LC864621 | 3a | - |
| 18 | 669 | *G. dehaani* | Iwate | Ichinoseki, Genbi | Yamaguchi | DA | LC864622 | 3a | DRR641607 |
| 19 | 670 | *G. dehaani* | Iwate | Ichinoseki, Genbi | Yamaguchi | DA | LC864623 | 3a | DRR641608 |
| 20 | 709 | *G. dehaani* | Miyagi | Ohira | Ouri | DA | LC864624 | 3a | - |
| 21 | 720 | *G. dehaani* | Miyagi | Taiwa | Yoshida | DA | LC864625 | 3a | - |
| 22 | 721 | *G. dehaani* | Miyagi | Taiwa | Yoshida | DA | - | - | DRR641618 |
| 23 | 689 | *G. dehaani* | Akita | Akita | Taiheihatta-odaishita | S | LC864626 | 3a | DRR641613 |
| 24 | 690 | *G. dehaani* | Akita | Akita | Taiheihatta-odaishita | S | LC864627 | 3a | DRR641614 |
| 25 | 727 | *G. dehaani* | Yamagata | Mamurogawa | Sasunabe | DA | LC864628 | 3a | DRR641619 |
| 26 | 728 | *G. dehaani* | Yamagata | Mamurogawa | Sasunabe | DA | LC864629 | 3a | DRR641620 |
| 27 | 674 | *G. dehaani* | Yamagata | Nanyo | Miyauchi | DA | LC864630 | 3a | DRR641609 |
| 28 | 675 | *G. dehaani* | Yamagata | Nanyo | Miyauchi | DA | LC864631 | 3a | DRR641610 |

| | | | | | | | | | |
|---|---|---|---|---|---|---|---|---|---|
| 29 | 755 | *G. dehaani* | Fukushima | Soma | Kamishikizawa | DA | LC864632 | 3a | - |
| 30 | 756 | *G. dehaani* | Fukushima | Soma | Kamishikizawa | DA | LC864633 | 3a | - |
| 31 | 699 | *G. dehaani* | Fukushima | Aizuwakamatsu, Ikki | Yahatasakashita | DA | LC864634 | 3a | DRR641615 |
| 32 | 700 | *G. dehaani* | Fukushima | Aizuwakamatsu, Ikki | Yahatasakashita | DA | LC864635 | 3a | DRR641616 |
| 33 | 825 | *G. dehaani* | Ibaraki | Hitachiota | Satogawa | un | LC864636 | 3a | - |
| 34 | 826 | *G. dehaani* | Ibaraki | Hitachiota | Satogawa | un | LC864637 | 3a | - |
| 35 | 922 | *G. dehaani* | Ibaraki | Sakuragawa, Makabe | Hatori | DA | LC864638 | 3a | DRR641692 |
| 36 | 923 | *G. dehaani* | Ibaraki | Sakuragawa, Makabe | Hatori | DA | LC864639 | 3a | - |
| 37 | 959 | *G. dehaani* | Tochigi | Ashikaga | Tsukiya | RE | LC864640 | 3a | - |
| 38 | 960 | *G. dehaani* | Tochigi | Ashikaga | Tsukiya | RE | LC864641 | 3a | - |
| 39 | 926 | *G. dehaani* | Tochigi | Nasukarasuyama | Kogisu | DA | LC864642 | 3a | DRR641693 |
| 40 | 927 | *G. dehaani* | Tochigi | Nasukarasuyama | Kogisu | DA | LC864643 | 3a | DRR641694 |
| 41 | 961 | *G. dehaani* | Gunma | Minakami | Tsukiyono | DA | LC864644 | 3a | - |
| 42 | 962 | *G. dehaani* | Gunma | Minakami | Tsukiyono | S | LC864645 | 3a | - |
| 43 | 982 | *G. dehaani* | Gunma | Minakami | Tsukiyono | S | LC864646 | 3a | - |
| 44 | 965 | *G. dehaani* | Gunma | Takasaki, Kurabuchi | Mizunuma | DA | LC864647 | 3a | DRR641707 |
| 45 | 967 | *G. dehaani* | Gunma | Takasaki, Kurabuchi | Mizunuma | DA | LC864648 | 3a | DRR641708 |
| 46 | 963 | *G. dehaani* | Saitama | Kamikawa | Shimoaguhara | DA | LC864649 | 3a | - |
| 47 | 964 | *G. dehaani* | Saitama | Kamikawa | Shimoaguhara | DA | LC864650 | 3a | DRR641706 |
| 48 | 983 | *G. dehaani* | Saitama | Kamikawa | Shimoaguhara | S | LC864651 | 3a | - |
| 49 | 1001 | *G. dehaani* | Chiba | Nagareyama | Kirigaya | BL | LC864652 | 3h | - |
| 50 | 1002 | *G. dehaani* | Chiba | Nagareyama | Kirigaya | BL | LC864653 | 3h | - |
| 51 | 998 | *G. dehaani* | Chiba | Narita | Otake | BL | LC864654 | 3h | DRR641712 |
| 52 | 999 | *G. dehaani* | Chiba | Narita | Otake | BL | LC864655 | 3h | DRR641713 |
| 53 | 1000 | *G. dehaani* | Chiba | Narita | Otake | BL | LC864656 | 3h | - |
| 54 | 1003 | *G. dehaani* | Chiba | Narita | Tsuchimuro | S | LC864657 | 3h | - |
| 55 | 1004 | *G. dehaani* | Chiba | Narita | Tsuchimuro | S | LC864658 | 3h | - |
| 56 | 892 | *G. dehaani* | Chiba | Tateyama | Koyatsu | BL | LC864659 | 3h | DRR641680 |
| 57 | 893 | *G. dehaani* | Chiba | Tateyama | Koyatsu | BL | LC864660 | 3h | DRR641681 |
| 58 | 894 | *G. dehaani* | Chiba | Tateyama | Koyatsu | BL | LC864661 | 3h | - |
| 59 | 895 | *G. dehaani* | Chiba | Tateyama | Koyatsu | BL | LC864662 | 3h | - |

| | | | | | | | | | |
|---|---|---|---|---|---|---|---|---|---|
| 60 | 879 | *G. dehaani* | Tokyo | Hachioji | Minamiasakawa | DA | LC864663 | 3a | DRR641679 |
| 61 | 880 | *G. dehaani* | Tokyo | Hachioji | Minamiasakawa | DA | LC864664 | 3a | - |
| 62 | 876 | *G. dehaani* | Kanagawa | Yokosuka | Abekura | BL | LC864665 | 3h | DRR641677 |
| 63 | 877 | *G. dehaani* | Kanagawa | Yokosuka | Abekura | BL | LC864666 | 3h | DRR641678 |
| 64 | 878 | *G. dehaani* | Kanagawa | Yokosuka | Abekura | BL | LC864667 | 3h | - |
| 65 | 644 | *G. dehaani* | Kanagawa | Chigasaki | Tsutsumi | BL | LC864668 | 3h | - |
| 66 | 874 | *G. dehaani* | Kanagawa | Matsuda | Yadoriki | DA | LC864669 | 3a | - |
| 67 | 875 | *G. dehaani* | Kanagawa | Matsuda | Yadoriki | DA | LC864670 | 3a | - |
| 68 | 645 | *G. dehaani* | Kanagawa | Yamakita | Mukohara | DA | LC864671 | 3a | - |
| 69 | 653 | *G. dehaani* | Niigata | Sado | Nishimikawa | DA | LC864672 | 3a | DRR641604 |
| 70 | 654 | *G. dehaani* | Niigata | Sado | Nishimikawa | DA | LC864673 | 3a | DRR641605 |
| 71 | 655 | *G. dehaani* | Niigata | Sado | Nishimikawa | DA | LC864674 | 3a | - |
| 72 | 765 | *G. dehaani* | Niigata | Nagaoka | Suyoshi | DA | LC864675 | 3a | DRR641626 |
| 73 | 766 | *G. dehaani* | Niigata | Nagaoka | Suyoshi | DA | LC864676 | 3a | DRR641627 |
| 74 | 759 | *G. dehaani* | Niigata | Itoigawa | Mushikawa | DA | LC864677 | 3a | - |
| 75 | 760 | *G. dehaani* | Niigata | Itoigawa | Mushikawa | DA | LC864678 | 3a | DRR641623 |
| 76 | 938 | *G. dehaani* | Toyama | Tonami | Ikuridani | DA | LC864679 | 3a | DRR641698 |
| 77 | 939 | *G. dehaani* | Toyama | Tonami | Ikuridani | DA | LC864680 | 3a | DRR641699 |
| 78 | 831 | *G. dehaani* | Ishikawa | Noto | Kami | S | - | - | DRR641659 |
| 79 | 832 | *G. dehaani* | Ishikawa | Noto | Kami | S | LC864681 | 3a | DRR641660 |
| 80 | 771 | *G. dehaani* | Ishikawa | Hakusan | Hakusan | DA | LC864682 | 3a | - |
| 81 | 772 | *G. dehaani* | Ishikawa | Hakusan | Hakusan | DA | LC864683 | 3a | - |
| 82 | 761 | *G. dehaani* | Fukui | Echizan | Yanagimoto | DA | LC864684 | 3a | DRR641624 |
| 83 | 762 | *G. dehaani* | Fukui | Echizan | Yanagimoto | DA | LC864685 | 3a | DRR641625 |
| 84 | 1029 | *G. dehaani* | Fukui | Mihama | Shinzyo | DA | LC864686 | 3a | - |
| 85 | 1030 | *G. dehaani* | Fukui | Mihama | Shinzyo | S | LC864687 | 3a | - |
| 86 | 928 | *G. dehaani* | Yamanashi | Minami-alps | AsiyasuAntsu | DA | LC864688 | 3a | DRR641695 |
| 87 | 929 | *G. dehaani* | Yamanashi | Minami-alps | AsiyasuAntsu | DA | LC864689 | 3a | - |
| 88 | 769 | *G. dehaani* | Nagano | Kizimadaira | Ogou | DA | LC864690 | 3a | - |
| 89 | 770 | *G. dehaani* | Nagano | Kizimadaira | Ogou | DA | LC864691 | 3a | - |
| 90 | 773 | *G. dehaani* | Nagano | Chikuma | Kuwabara | DA | LC864692 | 3a | DRR641629 |

| | | | | | | | | | |
|---|---|---|---|---|---|---|---|---|---|
| 91 | 774 | *G. dehaani* | Nagano | Chikuma | Kuwabara | DA | LC864693 | 3a | DRR641630 |
| 92 | 763 | *G. dehaani* | Nagano | Koumi | Chiyosato | DA | LC864694 | 3a | - |
| 93 | 764 | *G. dehaani* | Nagano | Koumi | Chiyosato | DA | LC864695 | 3a | - |
| 94 | 767 | *G. dehaani* | Nagano | Komagane | Akaho | DA | LC864696 | 3a | DRR641628 |
| 95 | 768 | *G. dehaani* | Nagano | Komagane | Akaho | DA | LC864697 | 3a | - |
| 96 | 415 | *G. dehaani* | Gifu | Minokamo, Miwa | Kawaura | DA | LC864698 | 3a | DRR641582 |
| 97 | 416 | *G. dehaani* | Gifu | Minokamo, Miwa | Kawaura | DA | LC864699 | 3a | - |
| 98 | 646 | *G. dehaani* | Shizuoka | Atami | Shimotaga | BL | LC864700 | 3h | - |
| 99 | 881 | *G. dehaani* | Shizuoka | Izu | Yoichizaka | BL | LC864701 | 3h | - |
| 100 | 882 | *G. dehaani* | Shizuoka | Izu | Yoichizaka | BL | LC864702 | 3h | - |
| 101 | 883 | *G. dehaani* | Shizuoka | Izu | Yoichizaka | BL | LC864703 | 3h | - |
| 102 | 884 | *G. dehaani* | Shizuoka | Izu | Yoichizaka | DA | LC864704 | 3h | - |
| 103 | 866 | *G. dehaani* | Shizuoka | Numazu | Ashitaka | BL | LC864705 | 3h | - |
| 104 | 867 | *G. dehaani* | Shizuoka | Numazu | Ashitaka | BL | LC864706 | 3h | DRR641675 |
| 105 | 868 | *G. dehaani* | Shizuoka | Numazu | Ashitaka | BL | - | - | DRR641676 |
| 106 | 869 | *G. dehaani* | Shizuoka | Numazu | Ashitaka | BL | LC864707 | 3h | - |
| 107 | 872 | *G. dehaani* | Shizuoka | Fujinomiya | Saori | DA | LC864708 | 3a | - |
| 108 | 873 | *G. dehaani* | Shizuoka | Fujinomiya | Saori | DA | LC864709 | 3a | - |
| 109 | 643 | *G. dehaani* | Shizuoka | Shizuoka, Shimizu | Shishihara | DA | LC864710 | 3a | - |
| 110 | 870 | *G. dehaani* | Shizuoka | Shizuoka, Shimizu | Shishihara | DA | LC864711 | 3a | - |
| 111 | 871 | *G. dehaani* | Shizuoka | Shizuoka, Shimizu | Shishihara | DA | LC864712 | 3a | - |
| 112 | 642 | *G. dehaani* | Shizuoka | Shimada | Oshiro | RE | LC864713 | 3a | - |
| 113 | 864 | *G. dehaani* | Shizuoka | Hamamatsu, Tenryu | Yamahigashi | DA | LC864714 | 3a | DRR641673 |
| 114 | 865 | *G. dehaani* | Shizuoka | Hamamatsu, Tenryu | Yamahigashi | DA | LC864715 | 3a | DRR641674 |
| 115 | 407 | *G. dehaani* | Aichi | Toyokawa | Toyosawa-tarumi | DA | LC864716 | 3a | - |
| 116 | 408 | *G. dehaani* | Aichi | Toyokawa | Toyosawa-tarumi | DA | LC864717 | 3a | - |
| 117 | 409 | *G. dehaani* | Aichi | Toyokawa | Toyosawa-tarumi | DA | LC864718 | 3a | - |
| 118 | 410 | *G. dehaani* | Aichi | Toyokawa | Toyosawa-tarumi | S | LC864719 | 3a | - |
| 119 | 411 | *G. dehaani* | Aichi | Toyokawa | Toyosawa-tarumi | S | LC864720 | 3a | - |
| 120 | 401 | *G. dehaani* | Aichi | Tahara, Ikawazu | Nagusa | DA | LC864721 | 3a | - |
| 121 | 402 | *G. dehaani* | Aichi | Tahara, Ikawazu | Nagusa | DA | LC864722 | 3a | - |

| | | | | | | | | | |
|---|---|---|---|---|---|---|---|---|---|
| 122 | 403 | *G. dehaani* | Aichi | Tahara, Ikawazu | Nagusa | RE | LC864723 | 3a | - |
| 123 | 404 | *G. dehaani* | Aichi | Tahara, Ikawazu | Nagusa | S | LC864724 | 3a | - |
| 124 | 405 | *G. dehaani* | Aichi | Tahara, Ikawazu | Nagusa | S | LC864725 | 3a | - |
| 125 | 412 | *G. dehaani* | Aichi | Toyota, Sanage | Hora | DA | LC864726 | 3a | - |
| 126 | 413 | *G. dehaani* | Aichi | Toyota, Sanage | Hora | DA | LC864727 | 3a | - |
| 127 | 414 | *G. dehaani* | Aichi | Toyota, Sanage | Hora | RE | LC864728 | 3a | - |
| 128 | 419 | *G. dehaani* | Aichi | Minamichita | Yamamiyugo | S | LC864729 | 3a | DRR641583 |
| 129 | 420 | *G. dehaani* | Aichi | Minamichita | Yamamiyugo | S | LC864730 | 3a | DRR641584 |
| 130 | 1005 | *G. dehaani* | Mie | Komono | Kamori | DA | LC864731 | 3a | - |
| 131 | 1006 | *G. dehaani* | Mie | Komono | Kamori | DA | LC864732 | 3a | - |
| 132 | 1009 | *G. dehaani* | Mie | Iga | Okubano | RE | LC864733 | 3a | - |
| 133 | 1010 | *G. dehaani* | Mie | Iga | Okubano | DA | LC864734 | 3a | - |
| 134 | 145 | *G. dehaani* | Mie | Ise | Yokowa | RE | LC864735 | 3a | DRR641574 |
| 135 | 341 | *G. dehaani* | Mie | Iitaka, Matsuzaka | Akaoke | un | LC864736 | 3a | - |
| 136 | 234 | *G. dehaani* | Mie | Watarai | Wakide | RE | LC864737 | 3a | - |
| 137 | 153 | *G. dehaani* | Mie | Shima, Ago | Ugata | RE | LC864738 | 3a | DRR641575 |
| 138 | 201 | *G. dehaani* | Mie | Taiki | Kamikusu | RE | LC864739 | 3a | - |
| 139 | 185 | *G. dehaani* | Mie | Taiki | Mayumi | DA | LC864740 | 3a | - |
| 140 | 270 | *G. dehaani* | Mie | Kihoku | Choshi riv.① | BL | LC864741 | 1b | - |
| 141 | 271 | *G. dehaani* | Mie | Kihoku | Choshi riv.① | DA | LC864742 | 1b | - |
| 142 | 272 | *G. dehaani* | Mie | Kihoku | Choshi riv.① | BL | LC864743 | 1b | - |
| 143 | 274 | *G. dehaani* | Mie | Kihoku | Choshi riv.② | BL | LC864744 | 1b | - |
| 144 | 275 | *G. dehaani* | Mie | Kihoku | Choshi riv.② | BL | LC864745 | 1b | - |
| 145 | 276 | *G. dehaani* | Mie | Kihoku | Choshi riv.③ | RE | LC864746 | 1b | DRR641578 |
| 146 | 277 | *G. dehaani* | Mie | Kihoku | Choshi riv.③ | BL | LC864747 | 1b | - |
| 147 | 278 | *G. dehaani* | Mie | Kihoku | Choshi riv.③ | RE | LC864748 | 1b | - |
| 148 | 279 | *G. dehaani* | Mie | Kihoku | Choshi riv.③ | BL | LC864749 | 1b | DRR641579 |
| 149 | 280 | *G. dehaani* | Mie | Kihoku | Choshi riv.④ | BL | LC864750 | 1b | - |
| 150 | 281 | *G. dehaani* | Mie | Kihoku | Choshi riv.④ | S | LC864751 | 1b | - |
| 151 | 282 | *G. dehaani* | Mie | Kihoku | Choshi riv.⑤ | RE | LC864752 | 1b | - |
| 152 | 283 | *G. dehaani* | Mie | Kihoku | Choshi riv.⑤ | S | LC864753 | 1b | - |

| | | | | | | | | | |
|---|---|---|---|---|---|---|---|---|---|
| 153 | 284 | *G. dehaani* | Mie | Kihoku | Choshi riv.⑤ | S | LC864754 | 1b | - |
| 154 | 285 | *G. dehaani* | Mie | Kihoku | Choshi riv.⑥ | DA | LC864755 | 1b | - |
| 155 | 125 | *G. dehaani* | Mie | Kihoku | Kawauchi | RE | LC864756 | 3a | - |
| 156 | 126 | *G. dehaani* | Mie | Kihoku | Kawauchi | DA | LC864757 | 3a | - |
| 157 | 164 | *G. dehaani* | Mie | Kihoku | Kouchi | DA | LC864758 | 3a | - |
| 158 | 265 | *G. dehaani* | Mie | Owase | Minamiura | BL | LC864759 | 1b | - |
| 159 | 239 | *G. dehaani* | Mie | Owase | Kada | BL | LC864760 | 1b | - |
| 160 | 213 | *G. dehaani* | Mie | Kumano, Ikusei | Ogawa | DA | LC864761 | 3a | - |
| 161 | 225 | *G. dehaani* | Mie | Kumano | Ido | BL | LC864762 | 1b | - |
| 162 | 860 | *G. dehaani* | Mie | Kumano, Arima | Ubuta riv. | BL | LC864763 | 1b | - |
| 163 | 861 | *G. dehaani* | Mie | Kumano, Arima | Ubuta riv. | BL | LC864764 | 1b | - |
| 164 | 862 | *G. dehaani* | Mie | Kumano, Arima | Ubuta riv. | BL | LC864765 | 1b | - |
| 165 | 863 | *G. dehaani* | Mie | Kumano, Arima | Ubuta riv. | RE | LC864766 | 1b | - |
| 166 | 189 | *G. dehaani* | Mie | Kiwa | Akagi | DA | LC864767 | 3a | DRR641576 |
| 167 | 207 | *G. dehaani* | Mie | Kiwa | Akagi | BL | LC864768 | 1b | DRR641577 |
| 168 | 208 | *G. dehaani* | Mie | Kiwa | Akagi | BL | LC864769 | 1b | - |
| 169 | 209 | *G. dehaani* | Mie | Kiwa | Akagi | DA | LC864770 | 3a | - |
| 170 | 101 | *G. dehaani* | Mie | Kumano, Kiwa | Senmaida① | BL | LC864771 | 1b | - |
| 171 | 136 | *G. dehaani* | Mie | Kumano, Kiwa | Senmaida② | BL | LC864772 | 1b | - |
| 172 | 137 | *G. dehaani* | Mie | Kumano, Kiwa | Senmaida② | BL | LC864773 | 1b | - |
| 173 | 843 | *G. dehaani* | Shiga | Sekigahara | Tama | DA | LC864774 | 3a | - |
| 174 | 844 | *G. dehaani* | Shiga | Sekigahara | Tama | DA | LC864775 | 3a | - |
| 175 | 794 | *G. dehaani* | Shiga | Takashima, Imazu | Tsunokawa | DA | LC864776 | 3a | - |
| 176 | 795 | *G. dehaani* | Shiga | Takashima, Imazu | Tsunokawa | DA | LC864777 | 3a | - |
| 177 | 1007 | *G. dehaani* | Shiga | Koka | Minakuchi | DA | LC864778 | 3a | DRR641714 |
| 178 | 1008 | *G. dehaani* | Shiga | Koka | Minakuchi | DA | - | - | DRR641715 |
| 179 | 817 | *G. dehaani* | Kyoto | Kyotango, Yasaka | Nonaka | RE | LC864779 | 3a | DRR641650 |
| 180 | 818 | *G. dehaani* | Kyoto | Kyotango, Yasaka | Nonaka | DA | LC864780 | 3a | DRR641651 |
| 181 | 819 | *G. dehaani* | Kyoto | Kyotango, Yasaka | Nonaka | S | LC864781 | 3a | - |
| 182 | 796 | *G. dehaani* | Kyoto | Kyoto, Sakyo | Ohara-kodeishi | DA | LC864782 | 3a | - |
| 183 | 797 | *G. dehaani* | Kyoto | Kyoto, Sakyo | Ohara-kodeishi | S | LC864783 | 3a | - |

| | | | | | | | | | |
|---|---|---|---|---|---|---|---|---|---|
| 184 | 816 | *G. dehaani* | Kyoto | Kizugawa, Yamashiro | Zindozi | DA | LC864784 | 3a | - |
| 185 | 798 | *G. dehaani* | Hyogo | Nishinomiya, Yamaguch | Funasaka | DA | LC864785 | 3a | - |
| 186 | 799 | *G. dehaani* | Hyogo | Nishinomiya, Yamaguch | Funasaka | DA | LC864786 | 3a | - |
| 187 | 627 | *G. dehaani* | Hyogo | Awaji | Notao | DA | LC864787 | 3a | DRR641599 |
| 188 | 628 | *G. dehaani* | Hyogo | Awaji | Notao | DA | LC864788 | 3a | DRR641600 |
| 189 | 821 | *G. dehaani* | Hyogo | Sayo | Sakurayama | RE | LC864789 | 3a | DRR641653 |
| 190 | 822 | *G. dehaani* | Hyogo | Sayo | Sakurayama | DA | LC864790 | 3a | - |
| 191 | 1011 | *G. dehaani* | Nara | Yamazoe | Zemyo | RE | LC864791 | 3a | - |
| 192 | 1012 | *G. dehaani* | Nara | Yamazoe | Zemyo | DA | LC864792 | 3a | - |
| 193 | 351 | *G. dehaani* | Nara | Kawakami | Takahara | un | LC864793 | 3a | - |
| 194 | 301 | *G. dehaani* | Nara | Gojo, Oto | Ui | DA | LC864794 | 3a | - |
| 195 | 302 | *G. dehaani* | Nara | Gojo, Oto | Ui | DA | LC864795 | 3a | - |
| 196 | 303 | *G. dehaani* | Nara | Gojo, Oto | Ui | S | LC864796 | 3a | - |
| 197 | 304 | *G. dehaani* | Nara | Gojo, Oto | Ui | DA | LC864797 | 3a | - |
| 198 | 305 | *G. dehaani* | Nara | Gojo, Oto | Ui | DA | LC864798 | 3a | - |
| 199 | 357 | *G. dehaani* | Nara | Nishiyoshino, Gojo | Kamino | DA | LC864799 | 3a | - |
| 200 | 116 | *G. dehaani* | Nara | Kamikitayama | Kawaai | DA | LC864800 | 1b | - |
| 201 | 254 | *G. dehaani* | Mie | Kamikitayama | Shirakawa | BL | LC864801 | 1b | - |
| 202 | 249 | *G. dehaani* | Nara | Shimokitayama | Shimoikehara | RE | LC864802 | 1b | - |
| 203 | 250 | *G. dehaani* | Nara | Shimokitayama | Shimoikehara | BL | LC864803 | 1b | - |
| 204 | 251 | *G. dehaani* | Nara | Shimokitayama | Shimoikehara | S | LC864804 | 1b | - |
| 205 | 306 | *G. dehaani* | Nara | Totsukawa | Asahi | un | LC864805 | 3a | - |
| 206 | 307 | *G. dehaani* | Nara | Totsukawa | Asahi | un | LC864806 | 3a | - |
| 207 | 308 | *G. dehaani* | Nara | Totsukawa | Asahi | un | LC864807 | 3a | - |
| 208 | 309 | *G. dehaani* | Nara | Totsukawa | Asahi | S | LC864808 | 3a | - |
| 209 | 310 | *G. dehaani* | Nara | Totsukawa | Asahi | S | LC864809 | 3a | - |
| 210 | 50 | *G. dehaani* | Nara | Totsukawa | Kamiyukawa | DA | LC864810 | 3a | - |
| 211 | 296 | *G. dehaani* | Nara | Totsukawa | Hiradani | DA | LC864811 | 3a | DRR641580 |
| 212 | 297 | *G. dehaani* | Nara | Totsukawa | Hiradani | DA | LC864812 | 3a | DRR641581 |
| 213 | 298 | *G. dehaani* | Nara | Totsukawa | Hiradani | DA | LC864813 | 3a | - |
| 214 | 299 | *G. dehaani* | Nara | Totsukawa | Hiradani | S | LC864814 | 3a | - |

| | | | | | | | | | |
|---|---|---|---|---|---|---|---|---|---|
| 215 | 300 | *G. dehaani* | Nara | Totsukawa | Hiradani | S | LC864815 | 3a | - |
| 216 | 244 | *G. dehaani* | Wakayama | Kitayama | Nanairo | DA | LC864816 | 3a | - |
| 217 | 245 | *G. dehaani* | Wakayama | Kitayama | Nanairo | DA | LC864817 | 3a | - |
| 218 | 246 | *G. dehaani* | Wakayama | Kitayama | Nanairo | DA | LC864818 | 3a | - |
| 219 | 247 | *G. dehaani* | Wakayama | Kitayama | Nanairo | DA | LC864819 | 3a | - |
| 220 | 248 | *G. dehaani* | Wakayama | Kitayama | Nanairo | S | LC864820 | 3a | - |
| 221 | 62 | *G. dehaani* | Wakayama | Kitayama | Onuma | DA | LC864821 | 1b | DRR641568 |
| 222 | 64 | *G. dehaani* | Wakayama | Kitayama | Onuma | BL | LC864822 | 1b | DRR641569 |
| 223 | 65 | *G. dehaani* | Wakayama | Kitayama | Onuma | BL | LC864823 | 1b | - |
| 224 | 9 | *G. dehaani* | Wakayama | Hashimoto | Hashiramoto | DA | LC864824 | 3a | - |
| 225 | 10 | *G. dehaani* | Wakayama | Hashimoto | Hashiramoto | DA | LC864825 | 3a | - |
| 226 | 11 | *G. dehaani* | Wakayama | Hashimoto | Hashiramoto | DA | LC864826 | 3a | - |
| 227 | 12 | *G. dehaani* | Wakayama | Hashimoto | Hashiramoto | DA | LC864827 | 3a | - |
| 228 | 13 | *G. dehaani* | Wakayama | Hashimoto | Hashiramoto | DA | LC864828 | 3a | - |
| 229 | 6 | *G. dehaani* | Wakayama | Wakayama | Takihata | DA | LC864829 | 3a | - |
| 230 | 7 | *G. dehaani* | Wakayama | Wakayama | Takihata | S | LC864830 | 3a | - |
| 231 | 1 | *G. dehaani* | Wakayama | Wakayama | Miyama | S | LC864831 | 3a | - |
| 232 | 2 | *G. dehaani* | Wakayama | Wakayama | Miyama | S | LC864832 | 3a | - |
| 233 | 3 | *G. dehaani* | Wakayama | Wakayama | Miyama | S | LC864833 | 3a | - |
| 234 | 4 | *G. dehaani* | Wakayama | Wakayama | Miyama | S | LC864834 | 3a | - |
| 235 | 5 | *G. dehaani* | Wakayama | Wakayama | Miyama | S | LC864835 | 3a | - |
| 236 | 14 | *G. dehaani* | Wakayama | Kudoyama | Kitamata | un | LC864836 | 3a | - |
| 237 | 29 | *G. dehaani* | Wakayama | Katsuragi | Hanazono | S | LC864837 | 3a | - |
| 238 | 30 | *G. dehaani* | Wakayama | Katsuragi | Hanazono | S | LC864838 | 3a | - |
| 239 | 31 | *G. dehaani* | Wakayama | Katsuragi | Hanazono | S | LC864839 | 3a | - |
| 240 | 32 | *G. dehaani* | Wakayama | Katsuragi | Hanazono | S | LC864840 | 3a | - |
| 241 | 33 | *G. dehaani* | Wakayama | Katsuragi | Hanazono | S | LC864841 | 3a | - |
| 242 | 34 | *G. dehaani* | Wakayama | Katsuragi | Hanazono | S | LC864842 | 3a | - |
| 243 | 28 | *G. dehaani* | Wakayama | Tanabe, Ryuzin | Mitsumata riv. | DA | LC864843 | 3a | - |
| 244 | 23 | *G. dehaani* | Wakayama | Tanabe, Ryuzin | Gomadan | DA | LC864844 | 3a | - |
| 245 | 24 | *G. dehaani* | Wakayama | Tanabe, Ryuzin | Gomadan | DA | LC864845 | 3a | - |

| | | | | | | | | | |
|---|---|---|---|---|---|---|---|---|---|
| 246 | 25 | *G. dehaani* | Wakayama | Tanabe, Ryuzin | Gomadan | S | LC864846 | 3a | - |
| 247 | 26 | *G. dehaani* | Wakayama | Tanabe, Ryuzin | Gomadan | S | LC864847 | 3a | - |
| 248 | 27 | *G. dehaani* | Wakayama | Tanabe, Ryuzin | Gomadan | S | LC864848 | 3a | - |
| 249 | 812 | *G. dehaani* | Wakayama | Kainan | Shimotsu | BL | LC864849 | 3a | - |
| 250 | 16 | *G. dehaani* | Wakayama | Kimino | Higashino | DA | LC864850 | 3a | - |
| 251 | 36 | *G. dehaani* | Wakayama | Hidaka | Shiga | DA | LC864851 | 3a | - |
| 252 | 38 | *G. dehaani* | Wakayama | Hidaka | Shiga | S | LC864852 | 3a | - |
| 253 | 53 | *G. dehaani* | Wakayama | Hidakagawa | Sansa | RE | LC864853 | 3a | - |
| 254 | 54 | *G. dehaani* | Wakayama | Hidakagawa | Sansa | RE | LC864854 | 3a | - |
| 255 | 68 | *G. dehaani* | Wakayama | Minabe | Nishihonzyo | DA | LC864855 | 3a | DRR641570 |
| 256 | 69 | *G. dehaani* | Wakayama | Minabe | Nishihonzyo | DA | LC864856 | 3a | DRR641571 |
| 257 | 70 | *G. dehaani* | Wakayama | Minabe | Nishihonzyo | DA | LC864857 | 3a | - |
| 258 | 71 | *G. dehaani* | Wakayama | Minabe | Nishihonzyo | S | LC864858 | 3a | - |
| 259 | 72 | *G. dehaani* | Wakayama | Minabe | Nishihonzyo | S | LC864859 | 3a | - |
| 260 | 73 | *G. dehaani* | Wakayama | Minabe | Nishihonzyo | S | LC864860 | 3a | - |
| 261 | 77 | *G. dehaani* | Wakayama | Kamitonda | Ikuma | RE | LC864861 | 3a | - |
| 262 | 78 | *G. dehaani* | Wakayama | Kamitonda | Ikuma | S | LC864862 | 3a | - |
| 263 | 96 | *G. dehaani* | Wakayama | Shirahama | Takegaito | S | LC864863 | 3a | - |
| 264 | 117 | *G. dehaani* | Wakayama | Tanabe | Ose | RE | LC864864 | 3a | - |
| 265 | 118 | *G. dehaani* | Wakayama | Tanabe | Ose | S | LC864865 | 3a | - |
| 266 | 119 | *G. dehaani* | Wakayama | Tanabe | Ose | RE | LC864866 | 3a | - |
| 267 | 120 | *G. dehaani* | Wakayama | Tanabe | Ose | DA | LC864867 | 3a | - |
| 268 | 121 | *G. dehaani* | Wakayama | Shingu | Kokonoe | DA | LC864868 | 3a | - |
| 269 | 122 | *G. dehaani* | Wakayama | Shingu | Kokonoe | DA | LC864869 | 3a | - |
| 270 | 123 | *G. dehaani* | Wakayama | Shingu | Kokonoe | - | LC864870 | 3a | - |
| 271 | 124 | *G. dehaani* | Wakayama | Shingu | Kokonoe | DA | LC864871 | 3a | - |
| 272 | 183 | *G. dehaani* | Wakayama | Shingu, Kumanogawa | Kuju | DA | LC864872 | 3a | - |
| 273 | 79 | *G. dehaani* | Wakayama | Nachikatsuura | Uragami | DA | LC864873 | 3a | DRR641572 |
| 274 | 80 | *G. dehaani* | Wakayama | Nachikatsuura | Uragami | RE | LC864874 | 3a | DRR641573 |
| 275 | 81 | *G. dehaani* | Wakayama | Nachikatsuura | Uragami | DA | LC864875 | 3a | - |
| 276 | 82 | *G. dehaani* | Wakayama | Nachikatsuura | Uragami | DA | LC864876 | 3a | - |

| | | | | | | | | | |
|---|---|---|---|---|---|---|---|---|---|
| 277 | 83 | *G. dehaani* | Wakayama | Nachikatsuura | Uragami | RE | LC864877 | 3a | - |
| 278 | 84 | *G. dehaani* | Wakayama | Nachikatsuura | Uragami | RE | LC864878 | 3a | - |
| 279 | 87 | *G. dehaani* | Wakayama | Susami | Esumi | RE | LC864879 | 3a | - |
| 280 | 1039 | *G. dehaani* | Wakayama | Kushimoto | Shionomisaki | RE | LC864880 | 3a | - |
| 281 | 829 | *G. dehaani* | Tottori | Wakasa | Mikura | DA | LC864881 | 3a | DRR641657 |
| 282 | 830 | *G. dehaani* | Tottori | Wakasa | Mikura | DA | LC864882 | 3a | DRR641658 |
| 283 | 833 | *G. dehaani* | Tottori | Yazu | Oe | DA | LC864883 | 3a | - |
| 284 | 834 | *G. dehaani* | Tottori | Yazu | Oe | DA | LC864884 | 3a | - |
| 285 | 814 | *G. dehaani* | Tottori | Daisen | Daisen | DA | LC864885 | 3b | - |
| 286 | 975 | *G. dehaani* | Tottori | Nanbu | Terauchi | BL | LC864886 | 3a | - |
| 287 | 996 | *G. dehaani* | Tottori | Nanbu | Terauchi | DA | LC864887 | 3a | - |
| 288 | 997 | *G. dehaani* | Tottori | Nanbu | Terauchi | DA | LC864888 | 3a | - |
| 289 | 800 | *G. dehaani* | Shimane | Okinoshima | Togo Isl., Harada | DA | LC864889 | 3b | DRR641644 |
| 290 | 801 | *G. dehaani* | Shimane | Okinoshima | Togo Isl., Harada | DA | LC864890 | 3b | DRR641645 |
| 291 | 839 | *G. dehaani* | Shimane | Nishinoshima | Nishinoshima Isl., Mita | DA | LC864891 | 3b | DRR641663 |
| 292 | 840 | *G. dehaani* | Shimane | Nishinoshima | Nishinoshima Isl., Mita | DA | LC864892 | 3b | DRR641664 |
| 293 | 793 | *G. dehaani* | Shimane | Matsue | Nishiikuma | DA | LC864893 | 3b | - |
| 294 | 813 | *G. dehaani* | Shimane | Okuizumo | Takezaki | S | LC864894 | 3a | - |
| 295 | 103 | *G. dehaani* | Shimane | Masuda, Hikimi | Hirose | un | LC864895 | 3b | - |
| 296 | 105 | *G. dehaani* | Shimane | Masuda, Hikimi | Hirose | S | LC864896 | 3b | - |
| 297 | 107 | *G. dehaani* | Shimane | Masuda, Hikimi | Hirose | S | LC864897 | 3b | - |
| 298 | 109 | *G. dehaani* | Shimane | Masuda, Hikimi | Hirose | S | LC864898 | 3b | - |
| 299 | 114 | *G. dehaani* | Shimane | Masuda, Hikimi | Hirose | S | LC864899 | 3b | - |
| 300 | 115 | *G. dehaani* | Shimane | Masuda, Hikimi | Hirose | un | LC864900 | 3b | - |
| 301 | 827 | *G. dehaani* | Okayama | Tsuyama | Yokoyama | RE | LC864901 | 3a | DRR641655 |
| 302 | 828 | *G. dehaani* | Okayama | Tsuyama | Yokoyama | RE | LC864902 | 3a | DRR641656 |
| 303 | 792 | *G. dehaani* | Okayama | Maniwa | Hiruzen-Shitao | DA | LC864903 | 3a | - |
| 304 | 815 | *G. dehaani* | Hiroshima | Syobara, Saizyo | Kumano | DA | LC864904 | 3b | - |
| 305 | 944 | *G. dehaani* | Hiroshima | Osakikamishima | Osakikamishima Isl., Higashino | DA | LC864905 | 3b | - |
| 306 | 945 | *G. dehaani* | Hiroshima | Osakikamishima | Osakikamishima Isl., Higashino | DA | LC864906 | 3b | - |
| 307 | 849 | *G. dehaani* | Hiroshima | Syobara, Saizyo | Kumano | DA | LC864907 | 3b | - |

| | | | | | | | | | |
|---|---|---|---|---|---|---|---|---|---|
| 308 | 850 | *G. dehaani* | Hiroshima | Syobara, Saizyo | Kumano | DA | LC864908 | 3b | - |
| 309 | 835 | *G. dehaani* | Hiroshima | Higashihiroshima | Saizyo | DA | LC864909 | 3b | - |
| 310 | 836 | *G. dehaani* | Hiroshima | Higashihiroshima | Saizyo | DA | LC864910 | 3b | - |
| 311 | 845 | *G. dehaani* | Hiroshima | Akiota | Ana | DA | LC864911 | 3b | DRR641667 |
| 312 | 846 | *G. dehaani* | Hiroshima | Akiota | Ana | DA | LC864912 | 3b | DRR641668 |
| 313 | 802 | *G. dehaani* | Yamaguchi | Hagi | Sasanami | BL | LC864913 | 3b | - |
| 314 | 803 | *G. dehaani* | Yamaguchi | Hagi | Sasanami | BL | LC864914 | 3b | - |
| 315 | 804 | *G. dehaani* | Yamaguchi | Hagi | Sasanami | BL | LC864915 | 3b | - |
| 316 | 820 | *G. dehaani* | Yamaguchi | Shimonoseki | Utsuikami | un | LC864916 | 3b | DRR641652 |
| 317 | 976 | *G. dehaani* | Yamaguchi | Shimonoseki | Utsuikami | S | LC864917 | 3b | - |
| 318 | 977 | *G. dehaani* | Yamaguchi | Shimonoseki | Utsuikami | S | LC864918 | 3b | - |
| 319 | 557 | *G. dehaani* | Tokushima | Naruto | Oshirotani riv. | DA | LC864919 | 3d | - |
| 320 | 560 | *G. dehaani* | Tokushima | Naruto | Oshirotani riv. | DA | LC864920 | 3d | - |
| 321 | 994 | *G. dehaani* | Tokushima | Miyoshi | Higashiiya-sugeoi | DA | LC864921 | 3b | - |
| 322 | 995 | *G. dehaani* | Tokushima | Miyoshi | Higashiiya-sugeoi | DA | LC864922 | 2b | - |
| 323 | 841 | *G. dehaani* | Tokushima | Miyoshi | Higashiiya-sugeoi | DA | LC864923 | 3b | DRR641665 |
| 324 | 842 | *G. dehaani* | Tokushima | Miyoshi | Higashiiya-sugeoi | DA | LC864924 | 2b | DRR641666 |
| 325 | 890 | *G. dehaani* | Tokushima | Miyoshi | Higashiiya-komi | BL | LC864925 | 1a | - |
| 326 | 568 | *G. dehaani* | Kagawa | Takamatsu | Goshikidai | DA | LC864926 | 3d | DRR641595 |
| 327 | 569 | *G. dehaani* | Kagawa | Takamatsu | Goshikidai | DA | LC864927 | 3d | DRR641596 |
| 328 | 570 | *G. dehaani* | Kagawa | Takamatsu | Goshikidai | DA | LC864928 | 3d | - |
| 329 | 572 | *G. dehaani* | Kagawa | Takamatsu | Goshikidai | DA | LC864929 | 3d | - |
| 330 | 901 | *G. dehaani* | Kagawa | Manno | Nakato | BL | LC864930 | 1a | - |
| 331 | 902 | *G. dehaani* | Kagawa | Manno | Nakato | BL | LC864931 | 1a | - |
| 332 | 903 | *G. dehaani* | Kagawa | Manno | Nakato | BL | LC864932 | 1a | - |
| 333 | 904 | *G. dehaani* | Kagawa | Manno | Nakato | BL | LC864933 | 1a | - |
| 334 | 466 | *G. dehaani* | Ehime | Matsuyama | Kukawa | DA | LC864934 | 3a | - |
| 335 | 467 | *G. dehaani* | Ehime | Matsuyama | Kukawa | RE | LC864935 | 3a | - |
| 336 | 990 | *G. dehaani* | Ehime | Matsuyama | Kugawa | S | LC864936 | 3a | - |
| 337 | 471 | *G. dehaani* | Ehime | Touon | Omogokei | DA | LC864937 | 3a | - |
| 338 | 472 | *G. dehaani* | Ehime | Touon | Omogokei | DA | LC864938 | 3a | - |

| | | | | | | | | | |
|---|---|---|---|---|---|---|---|---|---|
| 339 | 847 | *G. dehaani* | Ehime | Kumakogen | Wakayama | BL | LC864939 | 1a | DRR641669 |
| 340 | 848 | *G. dehaani* | Ehime | Kumakogen | Wakayama | BL | LC864940 | 1a | DRR641670 |
| 341 | 494 | *G. dehaani* | Ehime | Kumakogen | Nishitani | DA | LC864941 | 3a | - |
| 342 | 437 | *G. dehaani* | Ehime | Seiyo, Nomura | Matsutani | DA | LC864942 | 3a | DRR641587 |
| 343 | 441 | *G. dehaani* | Ehime | Seiyo, Nomura | Matsutani | DA | LC864943 | 3a | - |
| 344 | 946 | *G. dehaani* | Ehime | Ikata | Kojima | un | LC864944 | 3a | DRR641702 |
| 345 | 947 | *G. dehaani* | Ehime | Ikata | Kojima | S | LC864945 | 3a | DRR641703 |
| 346 | 431 | *G. dehaani* | Ehime | Uwajima | Tsushima | S | LC864946 | 1c | DRR641585 |
| 347 | 432 | *G. dehaani* | Ehime | Uwajima | Tsushima | RE | LC864947 | 1c | DRR641586 |
| 348 | 449 | *G. dehaani* | Kochi | Tosashimizu | Kagumi riv. | BL | LC864948 | 1c | - |
| 349 | 450 | *G. dehaani* | Kochi | Tosashimizu | Kagumi riv. | BL | LC864949 | 1c | - |
| 350 | 451 | *G. dehaani* | Kochi | Tosashimizu | Kagumi riv. | BL | LC864950 | 1c | - |
| 351 | 452 | *G. dehaani* | Kochi | Tosashimizu | Kagumi riv. | BL | LC864951 | 1c | - |
| 352 | 453 | *G. dehaani* | Kochi | Tosashimizu | Kagumi riv. | BL | LC864952 | 1c | - |
| 353 | 454 | *G. dehaani* | Kochi | Tosashimizu | Kagumi riv. | BL | LC864953 | 1c | - |
| 354 | 442 | *G. dehaani* | Kochi | Tosashimizu | Urajiri riv. | BL | - | - | DRR641588 |
| 355 | 443 | *G. dehaani* | Kochi | Tosashimizu | Urajiri riv. | BL | LC864954 | 1c | - |
| 356 | 444 | *G. dehaani* | Kochi | Tosashimizu | Urajiri riv. | BL | LC864955 | 1c | - |
| 357 | 446 | *G. dehaani* | Kochi | Tosashimizu | Urajiri riv. | S | LC864956 | 1c | DRR641589 |
| 358 | 447 | *G. dehaani* | Kochi | Tosashimizu | Urajiri riv. | S | LC864957 | 1c | - |
| 359 | 448 | *G. dehaani* | Kochi | Tosashimizu | Urajiri riv. | S | LC864958 | 1c | - |
| 360 | 886 | *G. dehaani* | Kochi | Otoyo | Sagayama | BL | LC864959 | 1a | - |
| 361 | 887 | *G. dehaani* | Kochi | Otoyo | Sagayama | BL | LC864960 | 1a | - |
| 362 | 888 | *G. dehaani* | Kochi | Otoyo | Sagayama | DA | LC864961 | 3a | - |
| 363 | 889 | *G. dehaani* | Kochi | Otoyo | Sagayama | DA | LC864962 | 3a | - |
| 364 | 584 | *G. dehaani* | Kochi | Muroto, Murotomisaki | Tsuro | BL | LC864963 | 1a | DRR641597 |
| 365 | 585 | *G. dehaani* | Kochi | Muroto, Murotomisaki | Tsuro | BL | LC864964 | 1a | DRR641598 |
| 366 | 508 | *G. dehaani* | Kochi | Kochi | Takagawa | BL | LC864965 | 1a | - |
| 367 | 511 | *G. dehaani* | Kochi | Kochi | Takagawa | DA | LC864966 | 2b | - |
| 368 | 524 | *G. dehaani* | Kochi | Kochi | Kagamiimai | BL | LC864967 | 2b | DRR641592 |
| 369 | 525 | *G. dehaani* | Kochi | Kochi | Kagamiimai | DA | LC864968 | 2b | - |

| | | | | | | | | | |
|---|---|---|---|---|---|---|---|---|---|
| 370 | 526 | *G. dehaani* | Kochi | Kochi | Kagamiimai | DA | LC864969 | 3b | - |
| 371 | 527 | *G. dehaani* | Kochi | Kochi | Kagamiimai | DA | LC864970 | 3b | DRR641593 |
| 372 | 991 | *G. dehaani* | Kochi | Kochi | Kagamiimai | DA | LC864971 | 3b | - |
| 373 | 498 | *G. dehaani* | Kochi | Ino | Kuwase | BL | LC864972 | 1a | DRR641590 |
| 374 | 499 | *G. dehaani* | Kochi | Ino | Kuwase | BL | LC864973 | 1a | - |
| 375 | 500 | *G. dehaani* | Kochi | Ino | Kuwase | DA | LC864974 | 3b | DRR641591 |
| 376 | 501 | *G. dehaani* | Kochi | Ino | Kuwase | RE | LC864975 | 3b | - |
| 377 | 482 | *G. dehaani* | Kochi | Tosa | Dema | RE | LC864976 | 3b | - |
| 378 | 486 | *G. dehaani* | Kochi | Tosa | Dema | RE | LC864977 | 3b | - |
| 379 | 487 | *G. dehaani* | Kochi | Suzaki | Kamibunhei | RE | LC864978 | 3b | - |
| 380 | 460 | *G. dehaani* | Kochi | Yusuhara | Nakahira | BL | LC864979 | 1c | - |
| 381 | 462 | *G. dehaani* | Kochi | Yusuhara | Nakahira | BL | LC864980 | 1c | - |
| 382 | 455 | *G. dehaani* | Kochi | Shimanto | Uchii riv. | RE | LC864981 | 1c | - |
| 383 | 459 | *G. dehaani* | Kochi | Shimanto | Uchii riv. | RE | LC864982 | 1c | - |
| 384 | 932 | *G. dehaani* | Fukuoka | itakyusyu, Kokuraminar | Yoshida | S | LC864983 | 2a | - |
| 385 | 933 | *G. dehaani* | Fukuoka | itakyusyu, Kokuraminar | Yoshida | S | LC864984 | 2b | - |
| 386 | 942 | *G. dehaani* | Fukuoka | Okagaki | Hatsu | RE | LC864985 | 2b | DRR641700 |
| 387 | 943 | *G. dehaani* | Fukuoka | Okagaki | Hatsu | DA | LC864986 | 3d | DRR641701 |
| 388 | 984 | *G. dehaani* | Fukuoka | Okagaki | Hatsu | DA | LC864987 | 2b | - |
| 389 | 985 | *G. dehaani* | Fukuoka | Okagaki | Hatsu | DA | LC864988 | 2b | - |
| 390 | 790 | *G. dehaani* | Fukuoka | Nogata | Kamitonno | DA | LC864989 | 2b | DRR641642 |
| 391 | 791 | *G. dehaani* | Fukuoka | Nogata | Kamitonno | DA | LC864990 | 2b | DRR641643 |
| 392 | 957 | *G. dehaani* | Fukuoka | Kurume, Yamamoto | Toyoda | DA | LC864991 | 2a | DRR641705 |
| 393 | 958 | *G. dehaani* | Fukuoka | Kurume, Yamamoto | Toyoda | DA | LC864992 | 2a | - |
| 394 | 934 | *G. dehaani* | Saga | Kanzaki, Sefuri | Haramaki | DA | LC864993 | 2a | - |
| 395 | 935 | *G. dehaani* | Saga | Kanzaki, Sefuri | Haramaki | S | LC864994 | 2a | - |
| 396 | 953 | *G. dehaani* | Saga | Shiraishi | Tsutsumi | RE | LC864995 | 3c | - |
| 397 | 954 | *G. dehaani* | Saga | Shiraishi | Tsutsumi | RE | LC864996 | 3c | - |
| 398 | 955 | *G. dehaani* | Saga | Tara | Ouraki | RE | LC864997 | 3c | - |
| 399 | 956 | *G. dehaani* | Saga | Tara | Ouraki | S | LC864998 | 2b | DRR641704 |
| 400 | 978 | *G. dehaani* | Saga | Tara | Ouraki | S | LC864999 | 3c | - |

| | | | | | | | | | |
|---|---|---|---|---|---|---|---|---|---|
| 401 | 1023 | *G. dehaani* | Nagasaki | Hirado, Ikitsuki | Ikitsukishima Isl., Minamimen | DA | LC865000 | 2b | - |
| 402 | 1024 | *G. dehaani* | Nagasaki | Hirado, Ikitsuki | Ikitsukishima Isl., Minamimen | DA | LC865001 | 2b | - |
| 403 | 1021 | *G. dehaani* | Nagasaki | Hirado | Hiradoshima Isl., Bougata | DA | LC865002 | 3c | DRR641720 |
| 404 | 1022 | *G. dehaani* | Nagasaki | Hirado | Hiradoshima Isl., Bougata | DA | LC865003 | 3c | DRR641721 |
| 405 | 1025 | *G. dehaani* | Nagasaki | Matsuura, Shisa | Takanomen | DA | LC865004 | 3c | - |
| 406 | 1026 | *G. dehaani* | Nagasaki | Matsuura, Shisa | Takanomen | DA | LC865005 | 3c | - |
| 407 | 930 | *G. dehaani* | Nagasaki | Nagayo | Yoshimutago | OC | LC865006 | 3c | DRR641696 |
| 408 | 931 | *G. dehaani* | Nagasaki | Nagayo | Yoshimutago | DA | LC865007 | 3c | - |
| 409 | 787 | *G. dehaani* | Nagasaki | Unzan, Agatsuma | Kawatokomyo | RE | LC865008 | 3c | DRR641640 |
| 410 | 788 | *G. dehaani* | Nagasaki | Unzan, Agatsuma | Kawatokomyo | OC | LC865009 | 3c | - |
| 411 | 789 | *G. dehaani* | Nagasaki | Unzan, Agatsuma | Kawatokomyo | OC | LC865010 | 3c | DRR641641 |
| 412 | 823 | *G. dehaani* | Nagasaki | Shinkamigoto | Nakadorijima Isl., Aokatago | DA | LC865011 | 3c | - |
| 413 | 824 | *G. dehaani* | Nagasaki | Shinkamigoto | Nakadorijima Isl., Aokatago | DA | LC865012 | 3c | DRR641654 |
| 414 | 980 | *G. dehaani* | Nagasaki | Goto, Kishikuma | Fukuejima Isl., Kawara | un | - | - | DRR641711 |
| 415 | 936 | *G. dehaani* | Nagasaki | Goto, Tamanoura | Fukuejima Isl., Arakawa | un | LC865013 | 3c | - |
| 416 | 937 | *G. dehaani* | Nagasaki | Goto, Tamanoura | Fukuejima Isl., Arakawa | un | LC865014 | 3c | DRR641697 |
| 417 | 661 | *G. dehaani* | Nagasaki | Goto | Danzyo, Oshima Isl. | un | LC865015 | 3i | DRR641606 |
| 418 | 1015 | *G. dehaani* | Kumamoto | Yamaga, Kahoku | Iwano | RE | LC865016 | 2b | - |
| 419 | 1016 | *G. dehaani* | Kumamoto | Yamaga, Kahoku | Iwano | DA | LC865017 | 2b | - |
| 420 | 1019 | *G. dehaani* | Kumamoto | Uto | Abiki | DA | LC865018 | 3d | - |
| 421 | 1020 | *G. dehaani* | Kumamoto | Uto | Abiki | S | LC865019 | 3d | - |
| 422 | 1017 | *G. dehaani* | Kumamoto | Yatsushiro, Kawata | Nishi | DA | LC865020 | 3d | DRR641718 |
| 423 | 1018 | *G. dehaani* | Kumamoto | Yatsushiro, Kawata | Nishi | DA | LC865021 | 3d | DRR641719 |
| 424 | 837 | *G. dehaani* | Kumamoto | Amakusa, Sumoto | Kamishima Isl., Kawachi | BL | LC865022 | 3c | DRR641661 |
| 425 | 838 | *G. dehaani* | Kumamoto | Amakusa, Sumoto | Kamishima Isl., Kawachi | RE | LC865023 | 3c | DRR641662 |
| 426 | 1013 | *G. dehaani* | Kumamoto | Amakusa | Gosyourajima Isl., Gosyoura | BL | LC865024 | 3c | DRR641716 |
| 427 | 1014 | *G. dehaani* | Kumamoto | Amakusa | Gosyourajima Isl., Gosyoura | OC | LC865025 | 3c | DRR641717 |
| 428 | 529 | *G. dehaani* | Kumamoto | Amakusa | Shimoshima Isl., Fukurengi | un | LC865026 | 3c | - |
| 429 | 530 | *G. dehaani* | Kumamoto | Amakusa | Shimoshima Isl., Fukurengi | un | LC865027 | 3c | DRR641594 |
| 430 | 981 | *G. dehaani* | Kumamoto | Amakusa | Shimoshima Isl., Takahamakita | BL | LC865028 | 3c | - |
| 431 | 810 | *G. dehaani* | Oita | Bungotakada | Matsuyuki, Nagaiwaya riv. | un | LC865029 | 2a | DRR641648 |

| | | | | | | | | | |
|---|---|---|---|---|---|---|---|---|---|
| 432 | 811 | *G. dehaani* | Oita | Bungotakada | Matsuyuki, Nagaiwaya riv. | un | LC865030 | 2a | DRR641649 |
| 433 | 650 | *G. dehaani* | Oita | Hita | Hidaka | DA | LC865031 | 2b | DRR641603 |
| 434 | 778 | *G. dehaani* | Oita | Hita | Hidaka | DA | LC865032 | 2b | DRR641632 |
| 435 | 986 | *G. dehaani* | Oita | Saiki | Kajiyoseura | OC | LC865033 | 3d | - |
| 436 | 987 | *G. dehaani* | Oita | Saiki | Kajiyoseura | OC | LC865034 | 3d | - |
| 437 | 896 | *G. dehaani* | Oita | Saiki | Kajiyoseura | OC | LC865035 | 3d | DRR641682 |
| 438 | 897 | *G. dehaani* | Oita | Saiki | Kajiyoseura | OC | LC865036 | 3b | DRR641683 |
| 439 | 940 | *G. dehaani* | Oita | Saiki | Ume, Minamitabaru | DA | LC865037 | 2b | - |
| 440 | 941 | *G. dehaani* | Oita | Saiki | Ume, Minamitabaru | DA | LC865038 | 3d | - |
| 441 | 988 | *G. dehaani* | Oita | Saiki | Ume, Minamitabaru | S | LC865039 | 2a | - |
| 442 | 989 | *G. dehaani* | Oita | Saiki | Ume, Minamitabaru | S | LC865040 | 2a | - |
| 443 | 808 | *G. dehaani* | Miyazaki | Nobeoka, Shimomiwa | Okita riv. | RE | LC865041 | 3d | - |
| 444 | 809 | *G. dehaani* | Miyazaki | Nobeoka, Shimomiwa | Okita riv. | RE | LC865042 | 3d | - |
| 445 | 783 | *G. dehaani* | Miyazaki | Gokase | Kuraoka | DA | LC865043 | 2a | DRR641637 |
| 446 | 784 | *G. dehaani* | Miyazaki | Gokase | Kuraoka | DA | LC865044 | 2a | DRR641638 |
| 447 | 805 | *G. dehaani* | Miyazaki | Kobayashi | Sukinakahara | DA | LC865045 | 2b | DRR641646 |
| 448 | 806 | *G. dehaani* | Miyazaki | Kobayashi | Sukinakahara | DA | LC865046 | 3d | DRR641647 |
| 449 | 807 | *G. dehaani* | Miyazaki | Kobayashi | Sukinakahara | DA | LC865047 | 3d | - |
| 450 | 979 | *G. dehaani* | Miyazaki | Kobayashi | Sukinakahara | DA | LC865048 | 3d | - |
| 451 | 851 | *G. dehaani* | Kagoshima | Nagashima | Nagashima Isl., Kawatoko | BL | LC865049 | 2b | DRR641671 |
| 452 | 852 | *G. dehaani* | Kagoshima | Nagashima | Nagashima Isl., Kawatoko | RE | LC865050 | 3d | DRR641672 |
| 453 | 992 | *G. dehaani* | Kagoshima | Nagashima | Nagashima Isl., Kawatoko | S | LC865051 | 3d | - |
| 454 | 993 | *G. dehaani* | Kagoshima | Nagashima | Nagashima Isl., Kawatoko | S | LC865052 | 3d | - |
| 455 | 924 | *G. dehaani* | Kagoshima | Satsumasendai, Iriki | Uranomyo | DA | LC865053 | 3d | - |
| 456 | 925 | *G. dehaani* | Kagoshima | Satsumasendai, Iriki | Uranomyo | RE | LC865054 | 3d | - |
| 457 | 779 | *G. dehaani* | Kagoshima | Ichikikushikino | Arakawa | RE | LC865055 | 3d | DRR641633 |
| 458 | 780 | *G. dehaani* | Kagoshima | Ichikikushikino | Arakawa | RE | LC865056 | 3d | DRR641634 |
| 459 | 951 | *G. dehaani* | Kagoshima | Hioki, Fukiage | Nagayoshi | BL | LC865057 | 3d | - |
| 460 | 952 | *G. dehaani* | Kagoshima | Hioki, Fukiage | Nagayoshi | S | LC865058 | 3d | - |
| 461 | 781 | *G. dehaani* | Kagoshima | Minamisatsuma | Kaseda-kominato | BL | LC865059 | 3d | DRR641635 |
| 462 | 782 | *G. dehaani* | Kagoshima | Minamisatsuma | Kaseda-kominato | BL | LC865060 | 3d | DRR641636 |

| | | | | | | | | | |
|---|---|---|---|---|---|---|---|---|---|
| 463 | 648 | *G. dehaani* | Kagoshima | Minamikyusyu, Ei | Kori | BL | LC865061 | 3d | DRR641602 |
| 464 | 776 | *G. dehaani* | Kagoshima | Minamikyusyu, Ei | Kori | BL | LC865062 | 3d | - |
| 465 | 777 | *G. dehaani* | Kagoshima | Shibushi | Noikura | DA | LC865063 | 2a | DRR641631 |
| 466 | 649 | *G. dehaani* | Kagoshima | Shibushi | Noikura | DA | LC865064 | 2a | - |
| 467 | 785 | *G. dehaani* | Kagoshima | Minamiosumi | Satahetsuka | BL | LC865065 | 3d | DRR641639 |
| 468 | 786 | *G. dehaani* | Kagoshima | Minamiosumi | Satahetsuka | BL | LC865066 | 2a | - |
| 469 | 647 | *G. dehaani* | Kagoshima | Minamiosumi | Satamagome | BL | LC865067 | 2b | DRR641601 |
| 470 | 775 | *G. dehaani* | Kagoshima | Minamiosumi | Satamagome | BL | LC865068 | 2b | - |
| 471 | 907 | *G. dehaani* | Kagoshima | Kamikoshiki | Kamikoshikijima Isl., Nakano | BL | LC865069 | 3g | DRR641686 |
| 472 | 908 | *G. dehaani* | Kagoshima | Kamikoshiki | Kamikoshikijima Isl., Nakano | BL | LC865070 | 3g | - |
| 473 | 909 | *G. dehaani* | Kagoshima | Kamikoshiki | Nakakoshikijima Isl., Taira | BL | LC865071 | 3g | - |
| 474 | 910 | *G. dehaani* | Kagoshima | Kamikoshiki | Nakakoshikijima Isl., Taira | BL | LC865072 | 3g | DRR641687 |
| 475 | 913 | *G. dehaani* | Kagoshima | Kamikoshiki | Nakakoshikijima Isl., Taira | BL | LC865073 | 3g | DRR641688 |
| 476 | 914 | *G. dehaani* | Kagoshima | Kamikoshiki | Nakakoshikijima Isl., Taira | BL | LC865074 | 3g | - |
| 477 | 916 | *G. dehaani* | Kagoshima | Shimokoshiki | Shimokoshikijima Isl., Aose | BL | LC865075 | 3g | DRR641689 |
| 478 | 915 | *G. dehaani* | Kagoshima | Shimokoshiki | Shimokoshikijima Isl., Katanoura | BL | LC865076 | 3g | - |
| 479 | 919 | *G. dehaani* | Kagoshima | Nishinoomote | Tanegashima Isl., Anzyo | BL | LC865077 | 3f | DRR641690 |
| 480 | 920 | *G. dehaani* | Kagoshima | Nishinoomote | Tanegashima Isl., Anzyo | S | LC865078 | 3f | DRR641691 |
| 481 | 921 | *G. dehaani* | Kagoshima | Nishinoomote | Tanegashima Isl., Anzyo | BL | LC865079 | 3f | - |
| 482 | 899 | *G. dehaani* | Kagoshima | Yakushima | Yakushima Isl., Anbo | BL | LC865080 | 3e | DRR641684 |
| 483 | 900 | *G. dehaani* | Kagoshima | Yakushima | Yakushima Isl., Anbo | BL | LC865081 | 3e | DRR641685 |
| 484 | 854 | *G. dehaani* | Kagoshima | Yakushima | Yakushima Isl., Kusukawa | BL | LC865082 | 3e | - |
| 485 | 855 | *G. dehaani* | Kagoshima | Yakushima | Yakushima Isl., Kusukawa | BL | LC865083 | 3e | - |
| 486 | 858 | *G. dehaani* | Kagoshima | Yakushima | Yakushima Isl., Koseda | RE | LC865084 | 3e | - |
| 487 | 856 | *G. dehaani* | Kagoshima | Yakushima | Yakushima Isl., Anbo | RE | LC865085 | 3e | - |
| 488 | 859 | *G. dehaani* | Kagoshima | Yakushima | Yakushima Isl., Mugio | BL | LC865086 | 3e | - |
| 489 | 857 | *G. dehaani* | Kagoshima | Yakushima | Yakushima Isl., Koshima | BL | LC865087 | 3e | - |
| 490 | 853 | *G. dehaani* | Kagoshima | Yakushima | Yakushima Isl., Nakama | BL | LC865088 | 3e | - |
| 491 | 973 | *G. dehaani* | Kagoshima | Toshima | Nakanoshima Isl. | S | LC865089 | 3e | - |
| 492 | 974 | *G. dehaani* | Kagoshima | Toshima | Nakanoshima Isl. | S | LC865090 | 3e | - |
| 493 | 651 | *G. exiguia* | Kagoshima | Minamiosumi | Satahetsuka | - | LC865091 | - | - |

| | | | | | | | | | |
|---|---|---|---|---|---|---|---|---|---|
| 494 | 652 | *G. exiguia* | Kagoshima | Minamiosumi | Satahetsuka | - | LC865092 | - | - |
| 495 | 905 | *G. koshikiensis* | Kagoshima | Kamikoshiki | Kamikoshikijima Isl., Nakano | - | LC865093 | - | - |
| 496 | 906 | *G. koshikiensis* | Kagoshima | Kamikoshiki | Kamikoshikijima Isl., Nakano | - | LC865094 | - | - |
| 497 | 911 | *G. koshikiensis* | Kagoshima | Kamikoshiki | Nakakoshikijima Isl., Taira | - | LC865095 | - | - |
| 498 | 912 | *G. koshikiensis* | Kagoshima | Kamikoshiki | Nakakoshikijima Isl., Taira | - | LC865096 | - | - |
| 499 | 917 | *G. koshikiensis* | Kagoshima | Shimokoshiki | Shimokoshikijima Isl., Aose | - | LC865097 | - | - |
| 500 | 918 | *G. koshikiensis* | Kagoshima | Shimokoshiki | Shimokoshikijima Isl., Aose | - | LC865098 | - | - |
| 501 | 885 | *G. marmorata* | Kagoshima | Yakushima | Yakushima Isl., Anbo | - | LC865099 | - | - |
| 502 | 898 | *G. marmorata* | Kagoshima | Yakushima | Yakushima Isl., Anbo | - | LC865100 | - | - |
| 503 | 949 | *G. sakamotoana* | Kagoshima | Tokunoshima | Tokunoshima Isl., Boma | - | LC865101 | - | - |
| 504 | 950 | *G. sakamotoana* | Okinawa | Naha | Okinawajima Isl., Syuritaira | - | LC865102 | - | - |
| - | - | *Geothelphusa* sp. | South Korea, Nampyung | Naju | - | - | MG674171 | 3c | - |
| - | - | *G. dehaani* | Tokyo | Hachizyo | Hachizyojima Isl., Nakanogo | - | LC743147 | 3a | - |
| - | - | *G. dehaani* | Tokyo | Hachizyo | Hachizyojima Isl., Nakanogo | - | LC743148 | 3a | - |
| - | - | *G. dehaani* | Kagoshima | Toshima | Kuchinoshima Isl. | - | LC743174 | 3e | - |
| - | - | *G. sakamotoana* | Kagoshima | Toshima | Takarajima Isl. | - | LC743302 | - | - |
| - | - | *G. sakamotoana* | Kagoshima | Toshima | Takarajima Isl. | - | LC743301 | - | - |
