## Supplementary material for "Genetic population structure of Japanese freshwater crab, *Geothelphusa dehaani* species complex (Decapoda: Potamidae) using genome-wide SNP": Table S2

Table S2.List of haplotypes. For sample numbers, refer to Table S1.

| Haplotypes. No. | Number of Haplotypes | Sample No. |
|---|---|---|
| Hap_1 | 20 | 1, 3, 5, 6, 7, 10, 11, 13, 14, 17, 19, 31, 37, 38, 44, 45, 92, 93, 224, 256 |
| Hap_2 | 4 | 99, 100, 101, 102 |
| Hap_3 | 4 | 98, 103, 104, 106 |
| Hap_4 | 2 | 77, 107 |
| Hap_5 | 4 | 86, 108, 109, 110 |
| Hap_6 | 1 | 111 |
| Hap_7 | 1 | 112 |
| Hap_8 | 56 | 23, 24, 72, 80, 81, 82, 83, 113, 114, 128, 129, 135, 136, 156, 160, 182, 195, 196, 197, 198, 199, 206, 208, 212, 213, 214, 217, 219, 229, 232, 233, 234, 236, 237, 238, 239, 240, 241, 242, 244, 245, 246, 247, 248, 249, 250, 251, 260, 263, 267, 268, 271, 342, 343, 362, 363 |
| Hap_9 | 3 | 115, 116, 118 |
| Hap_10 | 2 | 117, 119 |
| Hap_11 | 1 | 12 |
| Hap_12 | 3 | 120, 121, 122 |
| Hap_13 | 1 | 123 |
| Hap_14 | 1 | 124 |
| Hap_15 | 1 | 125 |
| Hap_16 | 10 | 96, 97, 126, 127, 133, 218, 220, 226, 230, 278 |
| Hap_17 | 1 | 130 |
| Hap_18 | 10 | 131, 132, 134, 138, 139, 176, 181, 187, 188, 192 |
| Hap_19 | 1 | 137 |
| Hap_20 | 7 | 140, 141, 148, 151, 154, 202, 204 |
| Hap_21 | 21 | 142, 144, 145, 146, 147, 149, 150, 152, 153, 159, 161, 163, 164, 167, 170, 171, 172, 200, 203, 221, 223 |
| Hap_22 | 1 | 143 |
| Hap_23 | 1 | 15 |
| Hap_24 | 1 | 155 |
| Hap_25 | 1 | 157 |
| Hap_26 | 1 | 158 |
| Hap_27 | 1 | 16 |
| Hap_28 | 3 | 162, 165, 168 |
| Hap_29 | 2 | 166, 270 |
| Hap_30 | 2 | 169, 194 |
| Hap_31 | 2 | 173, 174 |
| Hap_32 | 1 | 175 |
| Hap_33 | 6 | 177, 231, 235, 273, 276, 277 |

| | | |
|---|---|---|
| Hap_34 | 2 | 179, 281 |
| Hap_35 | 5 | 18, 27, 28, 30, 32 |
| Hap_36 | 1 | 180 |
| Hap_37 | 1 | 183 |
| Hap_38 | 1 | 184 |
| Hap_39 | 4 | 185, 186, 189, 190 |
| Hap_40 | 1 | 191 |
| Hap_41 | 4 | 193, 225, 227, 228 |
| Hap_42 | 1 | 2 |
| Hap_43 | 1 | 20 |
| Hap_44 | 1 | 201 |
| Hap_45 | 1 | 205 |
| Hap_46 | 2 | 207, 211 |
| Hap_47 | 3 | 209, 257, 258 |
| Hap_48 | 7 | 21, 33, 34, 35, 36, 39, 40 |
| Hap_49 | 4 | 210, 264, 265, 266 |
| Hap_50 | 1 | 215 |
| Hap_51 | 1 | 216 |
| Hap_52 | 1 | 222 |
| Hap_53 | 1 | 243 |
| Hap_54 | 2 | 25, 26 |
| Hap_55 | 1 | 252 |
| Hap_56 | 3 | 253, 254, 259 |
| Hap_57 | 1 | 255 |
| Hap_58 | 1 | 261 |
| Hap_59 | 1 | 262 |
| Hap_60 | 1 | 269 |
| Hap_61 | 1 | 272 |
| Hap_62 | 2 | 274, 275 |
| Hap_63 | 1 | 279 |
| Hap_64 | 1 | 280 |
| Hap_65 | 1 | 282 |
| Hap_66 | 7 | 283, 284, 286, 287, 288, 294, 303 |
| Hap_67 | 11 | 285, 293, 295, 297, 298, 299, 300, 304, 307, 308, 310 |
| Hap_68 | 4 | 289, 290, 291, 292 |
| Hap_69 | 1 | 29 |
| Hap_70 | 1 | 296 |
| Hap_71 | 1 | 301 |
| Hap_72 | 1 | 302 |

| | | |
|---|---|---|
| Hap_73 | 4 | 305, 309, 311, 312 |
| Hap_74 | 1 | 306 |
| Hap_75 | 3 | 313, 314, 315 |
| Hap_76 | 2 | 316, 318 |
| Hap_77 | 1 | 317 |
| Hap_78 | 6 | 319, 320, 326, 327, 328, 329 |
| Hap_79 | 2 | 321, 323 |
| Hap_80 | 3 | 322, 324, 369 |
| Hap_81 | 1 | 325 |
| Hap_82 | 1 | 330 |
| Hap_83 | 8 | 331, 332, 333, 339, 340, 360, 361, 366 |
| Hap_84 | 8 | 334, 335, 337, 338, 344, 345, LC743147, LC743148 |
| Hap_85 | 1 | 336 |
| Hap_86 | 1 | 341 |
| Hap_87 | 1 | 346 |
| Hap_88 | 1 | 347 |
| Hap_89 | 5 | 348, 349, 350, 351, 353 |
| Hap_90 | 1 | 352 |
| Hap_91 | 1 | 355 |
| Hap_92 | 1 | 356 |
| Hap_93 | 1 | 357 |
| Hap_94 | 1 | 358 |
| Hap_95 | 1 | 359 |
| Hap_96 | 2 | 364, 365 |
| Hap_97 | 1 | 367 |
| Hap_98 | 1 | 368 |
| Hap_99 | 3 | 370, 372, 377 |
| Hap_100 | 1 | 371 |
| Hap_101 | 2 | 373, 374 |
| Hap_102 | 2 | 375, 376 |
| Hap_103 | 1 | 378 |
| Hap_104 | 1 | 379 |
| Hap_105 | 1 | 380 |
| Hap_106 | 3 | 381, 382, 383 |
| Hap_107 | 2 | 384, 446 |
| Hap_108 | 3 | 385, 401, 402 |
| Hap_109 | 3 | 386, 389, 418 |
| Hap_110 | 1 | 387 |
| Hap_111 | 1 | 388 |

| | | |
|---|---|---|
| Hap_112 | 3 | 390, 391, 434 |
| Hap_113 | 5 | 392, 393, 395, 431, 432 |
| Hap_114 | 1 | 394 |
| Hap_115 | 10 | 396, 398, 403, 404, 405, 406, 408, 409, 410, 411 |
| Hap_116 | 1 | 397 |
| Hap_117 | 1 | 399 |
| Hap_118 | 1 | 400 |
| Hap_119 | 1 | 407 |
| Hap_120 | 2 | 41, 43 |
| Hap_121 | 2 | 412, 416 |
| Hap_122 | 1 | 413 |
| Hap_123 | 1 | 415 |
| Hap_124 | 1 | 417 |
| Hap_125 | 1 | 419 |
| Hap_126 | 1 | 42 |
| Hap_127 | 1 | 420 |
| Hap_128 | 1 | 421 |
| Hap_129 | 1 | 422 |
| Hap_130 | 1 | 423 |
| Hap_131 | 6 | 424, 427, 428, 429, 430, MG674171 |
| Hap_132 | 1 | 425 |
| Hap_133 | 1 | 426 |
| Hap_134 | 2 | 433, 439 |
| Hap_135 | 2 | 435, 437 |
| Hap_136 | 1 | 436 |
| Hap_137 | 1 | 438 |
| Hap_138 | 1 | 440 |
| Hap_139 | 4 | 441, 465, 466, 468 |
| Hap_140 | 1 | 442 |
| Hap_141 | 2 | 443, 444 |
| Hap_142 | 1 | 445 |
| Hap_143 | 1 | 447 |
| Hap_144 | 2 | 448, 449 |
| Hap_145 | 1 | 450 |
| Hap_146 | 1 | 451 |
| Hap_147 | 2 | 452, 453 |
| Hap_148 | 1 | 454 |
| Hap_149 | 2 | 455, 456 |
| Hap_150 | 1 | 457 |

| | | |
|---|---|---|
| Hap_151 | 1 | 458 |
| Hap_152 | 2 | 459, 460 |
| Hap_153 | 3 | 46, 47, 48 |
| Hap_154 | 1 | 461 |
| Hap_155 | 1 | 462 |
| Hap_156 | 2 | 463, 464 |
| Hap_157 | 1 | 467 |
| Hap_158 | 1 | 469 |
| Hap_159 | 1 | 470 |
| Hap_160 | 1 | 471 |
| Hap_161 | 1 | 472 |
| Hap_162 | 1 | 473 |
| Hap_163 | 1 | 474 |
| Hap_164 | 1 | 475 |
| Hap_165 | 1 | 476 |
| Hap_166 | 2 | 477, 478 |
| Hap_167 | 1 | 479 |
| Hap_168 | 2 | 480, 481 |
| Hap_169 | 4 | 482, 488, 489, 490 |
| Hap_170 | 2 | 483, 486 |
| Hap_171 | 1 | 484 |
| Hap_172 | 1 | 485 |
| Hap_173 | 1 | 487 |
| Hap_174 | 9 | 49, 50, 51, 52, 53, 54, 55, 56, 59 |
| Hap_175 | 1 | 491 |
| Hap_176 | 1 | 492 |
| Hap_177 | 1 | 57 |
| Hap_178 | 1 | 58 |
| Hap_179 | 1 | 60 |
| Hap_180 | 1 | 61 |
| Hap_181 | 4 | 62, 63, 64, 65 |
| Hap_182 | 2 | 66, 67 |
| Hap_183 | 1 | 68 |
| Hap_184 | 2 | 69, 71 |
| Hap_185 | 1 | 70 |
| Hap_186 | 5 | 73, 88, 89, 90, 91 |
| Hap_187 | 2 | 74, 75 |
| Hap_188 | 1 | 76 |
| Hap_189 | 1 | 79 |

| | | |
|---|---|---|
| Hap_190 | 2 | 8, 9 |
| Hap_191 | 1 | 84 |
| Hap_192 | 1 | 85 |
| Hap_193 | 2 | 87, 94 |
| Hap_194 | 1 | 95 |
| Hap_195 | 1 | LC743174 |
