## Supplementary material for "Genetic population structure of Japanese freshwater crab, *Geothelphusa dehaani* species complex (Decapoda: Potamidae) using genome-wide SNP": Table S3

Table S3. Number of haplotypes, Haplotype diversity (*h*), nucleotide diversity (π), Tajima's *D* and Fu's *Fs* for each clade based on mtDNA.

| | Number of individuals | Number of Haplotypes | *h* | π | Tajima's *D* | Tajima's *D* p-value | Fu's $F_S$ | $F_S$ p-value |
|---|---|---|---|---|---|---|---|---|
| 1 | 66 | 23 | 0.016671 +/- 0.008568 | 0.8709 +/- 0.0315 | -0.56156 | 0.326 | -0.96108 | 0.43055 |
| 2 | 37 | 20 | 0.027383 +/- 0.013878 | 0.9550 +/- 0.0157 | 0.66468 | 0.80295 | 0.05144 | 0.5522 |
| 3 | 386 | 152 | 0.023233 +/- 0.011566 | 0.9704 +/- 0.0049 | -1.68854 | 0.0098 | -23.66117 | 0.006 |
