## Supplementary material for "Genetic population structure of Japanese freshwater crab, *Geothelphusa dehaani* species complex (Decapoda: Potamidae) using genome-wide SNP": Table S4

Table S4. *F* st and $F_{ST}$ *p* -values among the five populations of SNPs.
Bottom left: Pairwise fixation index ($F_{ST}$), top right: $F_{ST}$ *p* -values.

| $F_{ST}$ \ *p* -value | 1 | 2 | 3 |
|---|---|---|---|
| 1 | | 0.00000+-0.0000 | 0.00000+-0.0000 |
| 2 | 0.73186 | | 0.00000+-0.0000 |
| 3 | 0.74908 | 0.75244 | |
