## Supplementary material for "Genetic population structure of Japanese freshwater crab, *Geothelphusa dehaani* species complex (Decapoda: Potamidae) using genome-wide SNP": Table S5

Table S5. In each ADMIXTURE analysis based on the SNPs data, sample size (N), the number of SNPs, and the CV error values.

| | N | number of SNPs | CV error | | | | | | | | | |
|---|---|---|---|---|---|---|---|---|---|---|---|---|
| | | | K=1 | K=2 | K=3 | K=4 | K=5 | K=6 | K=7 | K=8 | K=9 | K=10 |
| Set 0 | 154 | 308 | 0.37339 | 0.31692 | 0.30444 | 0.2898 | 0.28456 | 0.28543 | 0.28652 | 0.29713 | 0.30057 | 0.30672 |
| Set 1 | 17 | 595 | 1.41014 | 0.98298 | 1.25813 | 1.03843 | 0.82241 | 0.52991 | 0.54916 | 0.70742 | 0.42662 | 0.24137 |
| Set 2 | 16 | 820 | 1.44945 | 1.25993 | 1.60723 | 1.22194 | 1.03663 | 0.98336 | 0.93175 | 0.82748 | 0.56025 | 0.42699 |
| Set 3 | 16 | 856 | 1.54503 | 1.32577 | 1.46224 | 1.67223 | 1.34857 | 0.95495 | 0.9506 | 0.93307 | 0.49126 | 0.48682 |
| Set 4 | 21 | 236 | 1.28933 | 1.16137 | 1.32213 | 1.31876 | 1.47563 | 1.24735 | 1.0844 | 0.83889 | 0.62823 | 0.53658 |
| Set 5 | 31 | 590 | 0.9496 | 0.85144 | 0.91093 | 0.90639 | 0.96146 | 0.85026 | 0.8369 | 0.68702 | 0.87332 | 0.52633 |
| Set 6 | 39 | 410 | 0.87822 | 0.80712 | 0.84485 | 0.92308 | 0.91198 | 0.98354 | 0.92093 | 0.97935 | 0.81738 | 0.6914 |
