## Supplementary material for "Genetic population structure of Japanese freshwater crab, *Geothelphusa dehaani* species complex (Decapoda: Potamidae) using genome-wide SNP": Table S6

Table S6. Results of the scenario choice by DIYABC-RF analysis.

| | Body color | | | Area (Honshu vs. Kyushu) | | | | sKK first | | | | SHI first | | | |
|---|---|---|---|---|---|---|---|---|---|---|---|---|---|---|---|
| Scenario | S1 | S2 | S3 | S4 | S5 | S6 | S7 | S8 | S9 | S10 | S11 | S12 | S13 | S14 | S15 |
| Votes | 0 | 0 | 2 | 8 | 14 | 2 | 9 | 8 | 0 | 1 | 2 | 8 | 129 | 13 | 42 |

| | cK first | | | | nKC first | | | | HO first | | | | | multi divergence | | | |
|---|---|---|---|---|---|---|---|---|---|---|---|---|---|---|---|---|---|
| | S16 | S17 | S18 | S19 | S20 | S21 | S22 | S23 | S24 | S25 | S26 | S27 | S28 | S29 | S30 | S31 | S32 |
| | 1 | 3 | 22 | 3 | 1 | 11 | 7 | 5 | 0 | 17 | 3 | 3 | 3 | 38 | 520 | 62 | 63 |
