## Supplementary material for "Genetic population structure of Japanese freshwater crab, *Geothelphusa dehaani* species complex (Decapoda: Potamidae) using genome-wide SNP": Table S7

Table S7. Prior distribution parameters in DIYABC-RF analysis for five populations of the Sawagani (*Geothelphusa dehaani* species complex).

| Parameter | Minimum | Maximum |
|---|---|---|
| *Effective population size* | | |
| N1 (sKK population) | 10 | 100000 |
| N2 (SHI population) | 10 | 500000 |
| N3 (nKC population) | 10 | 100000 |
| N4 (cK population) | 10 | 100000 |
| N5 (HO population) | 10 | 500000 |
| *Time scale in generations* | | |
| t1 | 10 | 500000 |
| t2 | 10 | 500000 |
| t3 | 10 | 500000 |
| t4 | 10 | 500000 |
